# Structural basis for binding of RILPL1 to TMEM55B reveals a lysosomal platform for adaptor assembly through a conserved TBM motif

**DOI:** 10.1101/2025.08.19.670962

**Authors:** Dieter Waschbüsch, Prosenjit Pal, Raja S. Nirujogi, Melanie Cavin, Jaijeet Singh, Dario R. Alessi, Amir R. Khan

## Abstract

Inherited mutations in VPS35 and the kinase LRRK2 lead to hyperphosphorylation of Rab GTPases and promote the formation of phospho-Rab signalling complexes. A subset of RH2 domain-containing proteins from the RILP-homology family, including RILP, RILPL1, RILPL2, JIP3, and JIP4 are Rab effectors that recognize the LRRK2-phosphorylated switch 2 threonine of phospho-Rab8A and phospho-Rab10. More recently, phospho-Rabs have been found on lysosomal membranes within multi-protein assemblies involving TMEM55B and RILPL1. TMEM55B is a 284-residue lysosomal membrane protein with no homology to known proteins. It comprises a 218-residue cytosolic N-terminal region and two predicted transmembrane α-helices. Residues 80– 160, which face the cytosol, mediate binding to a C-terminal motif of RILPL1, formed after RILPL1 associates with phospho-Rab8A. Here, we report the crystal structures of TMEM55B alone and in complex with a C-terminal RILPL1 peptide, encompassing the TMEM55B interaction region, which we define as the TMEM55B Binding Motif (TBM). The cytosolic domain of TMEM55B adopts a rigid architecture of two tandem RING-like domains, each forming a Zn²⁺-stabilized 40-residue β-sandwich. TBM binding is mediated primarily by backbone hydrogen bonding and anchored by two glutamate residues from RILPL1. These findings support a model in which RILPL1 is recruited to phospho-Rab8A–positive lysosomes prior to TMEM55B engagement. Further co-immunoprecipitation and mutational analyses indicate that TMEM55B forms complexes independently of phospho-Rabs with proteins containing a conserved TBM, like that of RILPL1, including JIP3, JIP4, OCRL, WDR81, and TBC1D9B. Together, these findings uncover previously unrecognized regulatory networks associated with TMEM55B and lysosomal function and suggest that TMEM55B serves as a central hub for adaptor recruitment at the lysosomal membrane.

## INTRODUCTION

Parkinson’s disease (PD) is a disorder of the central nervous system that manifests as a progressive degeneration of motor mobility, loss of balance, and tremors. Features of the pathology include loss of dopaminergic neurons in the midbrain and the presence of protein aggregates termed Lewy bodies, composed mainly of α-synuclein, in surviving neurons^1^. About 10% of cases have a genetic basis, with mutations in the *Leucine-Rich Repeat Kinase 2* (*LRRK2) gene being among the most common* ^2^. The gene product is a 2,527-residue protein with multiple domains belonging to the ROCO family that is involved in regulation of autophagy, mitochondria, and Golgi dynamics^3^. LRRK2 pathogenic mutations such as R1441C are inherited in an autosomal dominant fashion and lead to hyperactive kinase activity. In addition to LRRK2, another common genetic inheritance is VPS35, which is a component of the retromer complex that transports endosomal cargo to Golgi and plasma membrane compartments. The D620N mutation in VPS35 is also autosomal dominant and leads to hyperactive LRRK2 kinase activity by an unknown mechanism.

LRRK2 substrates are a subset of Rab GTPases that includes Rab8A and Rab10, thereby connecting LRRK2 functions to membrane trafficking^4^. Rabs comprise the largest group (∼70 members) of the Ras superfamily, and they cycle between an active GTP-bound and inactive GDP form to regulate aspects of vesicle formation, transport, and organelle dynamics in eukaryotes^5^. The ‘switch’ regions of Rabs, termed switch 1 and 2, undergo local conformational changes that enable the recruitment of GTP-specific effectors to subcellular compartments, which subsequently control membrane trafficking through recruitment of effector proteins^6^. LRRK2 phosphorylates Rab8A at T72 and Rab10 at T73, conserved threonine residues located on the α-helical switch 2 region. This post-translational modification modulates interactions between Rabs and their binding partners^4,7^. We have previously characterized the molecular basis for phospho-Rab recruitment of effector proteins following LRRK2 phosphorylation of Rab8A. The structure of phospho-Rab8A (pRab8A) in complex with effector RILPL2 (Rab-interacting lysosome-associated protein-like 2) revealed a coiled-coil with an N-terminal X-shaped feature, termed an ‘X-cap’ that together encode both Rab-GTP and phospho-threonine (pT72) specificity^8^. This region of the effector protein is also known as the RILP homology domain 2 (RH2), and it is also found in RILP, RILPL1, JIP3 and JIP4.

In recent work, a proteomics approach identified a protein termed TMEM55B which interacts with pRab8A/RILPL1 complexes ^9^. TMEM55B is a 284-residue lysosomal protein with little sequence resemblance to other proteins and has been annotated as a phosphoinositol phosphatase^10^. From Alphafold3 predictions, the N-terminus of the protein (1-218) faces the cytosol, while the C-terminus consists of 2 transmembrane α-helices (219-239 and 249-269). TMEM55B lacks sequence similarities to other lipid phosphatases and the *in-vitro* activity showing dephosphorylation has not been verified in the literature^11^. Although TMEM55B has not generally been associated with Parkinson’s disease, lysosomal trafficking and autophagy are strongly linked to LRRK2 functions^12,13^.

Interactions between the RILPL1/TMEM55B complex have been localized to a globular segment 80-160 of TMEM55B that faces the cytosol^9^. The corresponding RILP1 interacting region, a cytosolic adaptor protein, comprises the C-terminal 12 residues (391-403) which lie within a highly flexible segment following the RH2 domain (residues 290-326). Here we have determined structures of the cytosolic globular domain of TMEM55B (80-160) both alone and in complex with a peptide segment 391-403 of RILPL1. The structures reveal a tandem module of two RING-type Zn^2+^-finger domains that resemble ubiquitin E3 ligases despite minimal sequence identities. The RILPL1 peptide interacts in an extended fashion along one face of the saucer-like TMEM55B through two glutamate anchors and a remarkable number of backbone hydrogen bonds. We designate this recognition sequence as the TMEM55B Binding Motif (TBM). We provide evidence that TMEM55B forms complexes independently of phospho-Rabs with a set of scaffolding proteins that contain a conserved TBM, like that of RILPL1, including OCRL, WDR81, JIP3, JIP4, and TBC1D9B. Together, these findings suggest that TMEM55B functions as a central hub for recruiting TBM-containing adaptor proteins to the lysosomal membrane.

## RESULTS

### Structure of TMEM55B resembles RING domains

The first ∼75 cytosolic residues of TMEM55B are predicted to be disordered, while the subsequent globular region 80-160 bears little sequence resemblance to known proteins in databases (**Figure 1A**). The X-ray structure of the segment 80-166 of TMEM55B was determined both in isolation and bound to a peptide encompassing residues 391-403 of RILPL1 (**Figure 1A**). This cytosolic segment of TMEM55B adopts two tandem β-sandwich domains comprising residues 88-116 and 126-158. Each of these two domains adopts a RING-like fold with a Zn^2+^ ion coordinated by four Cys residues in tetrahedral geometry (**Figure 1B**). The entire segment 86-166 comprising the two RING-like domains can be considered a single globular unit connected by a proline-rich linker (117-125). There are extensive polar and non-polar interactions between the two domains, including a salt bridge between Glu116 and Arg151. There are no significant conformational differences between TMEM55B alone and in complex with RILPL1, except for a flexible C-terminal tail that mediates slightly different crystal contacts (**Suppl Figure S1**).

**Figure 1:**
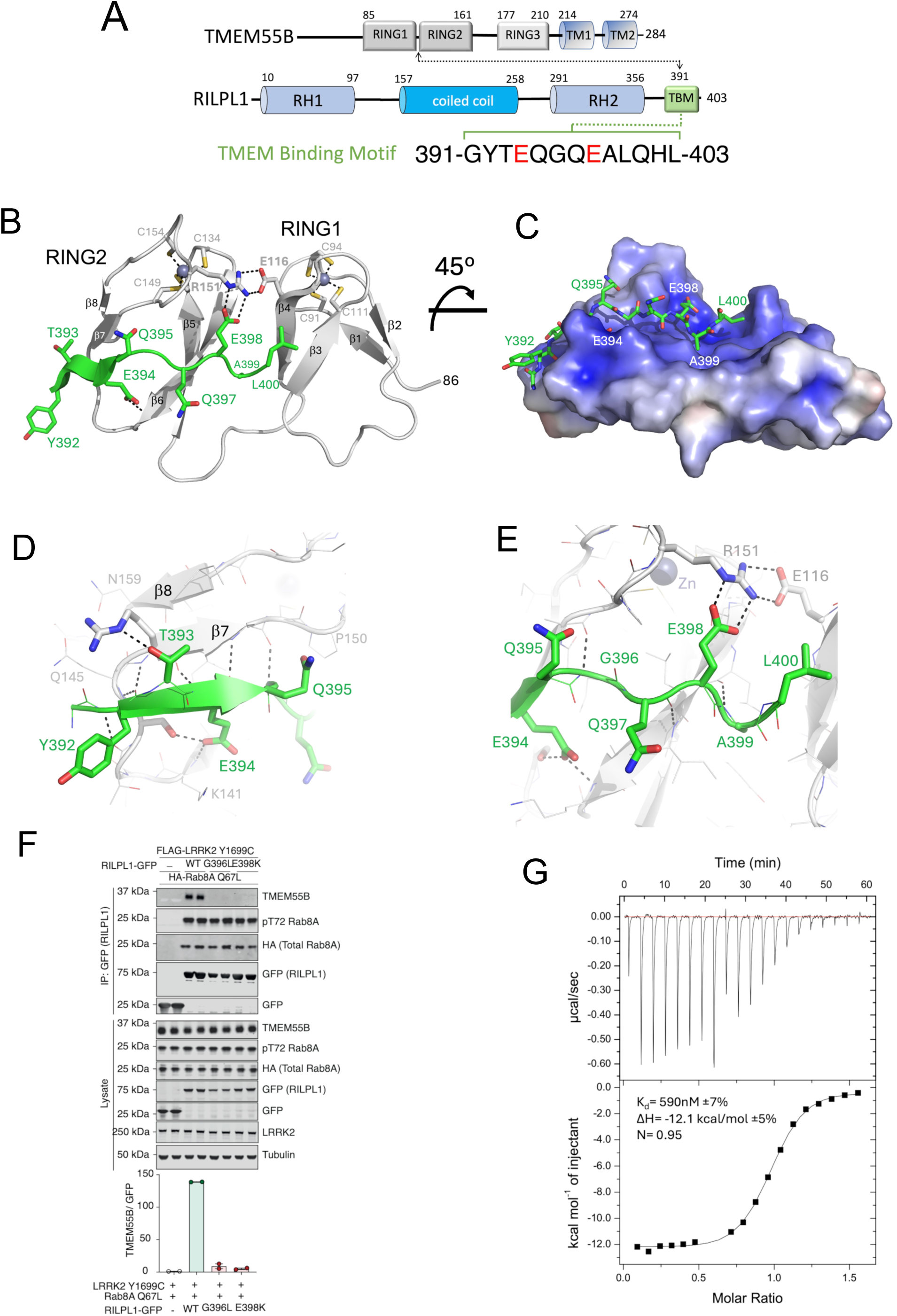
Structure of RILPL1 peptide in complex with the tandem RING-like domains of TMEM55B. **(A)** Domain organization of TMEM55B and RILPL1, with location of the TMEM binding motif (TBM). **(B)** Structure of the complex between RILPL1 (green) and TMEM55B (grey) in ribbon models. Key interacting residues are shown as stick models. **(C)** Surface electrostatic representation of TMEM55B with a stick model of the RILPL1 peptide. **(D)** Close-up of the interactions between the N-terminal part of RILPL1 peptide and TMEM55B, revealing the extent of backbone hydrogen bonds (dashed black lines). E398^RL1^ makes a key hydrogen bond with the backbone of K141^TM^. **(E)** Similarly, the C-terminal part of the peptide is shown, with the key salt bridge between E398^RL1^ and R151^TM^. **(F)** HEK293 TMEM55B/A double knock-out cells were transiently transfected with HA-Rab8A [Q67L] (GTP-bound mutant) and FLAG-LRRK2[Y1699C] (kinase-active mutant) either in the presence of RILPL1-GFP wild-type (WT) or G396L/E398K double mutant. GFP immunoprecipitation was performed and analysed by quantitative immunoblot analysis using the LI-COR Odyssey CLx Western blot imaging system and indicated antibodies. Quantitation of immuno-blotting data (performed using ImageStudioLite software version5.2.5, RRID:SCR_013715) is shown as mean ± SEM. **(G)** Isothermal titration calorimetry analysis of the interaction between RILPL1 peptide and TMEM55B. The peptide was injected into the ITC200 cell (Particular Sciences) containing TMEM55B as described in **Methods**.

### Structure of the complex between a RILPL1 peptide and TMEM55B

Descriptions of the complex will utilize ‘RL1’ and ‘TM’ superscripts as acronyms above residue numbers for RILPL1 and TMEM55B, respectively. RILPL1 (391-400) adopts an extended conformation that spreads across one face of saucer shaped TMEM55B (region 80-166), bridging both RING domains (**Figure 1B,C**). Residues 392-395 of RILPL1 form a parallel β-strand to complement a 2-stranded β-sheet in the C-terminal RING-like domain of TMEM55B. The backbone NH of Y392^RL1^ also makes a backbone H-bond with T143^TM^ (**Figure 1D**). Following this strand, G396^RL1^ marks a bend that directs residues 397-400 toward the N-terminal RING-like domain (**Figure 1E**). There are two additional backbone H-bonds between Q397^RL1^ and A399^RL1^ with β6 of the C-terminal RING-like domain. These intimate interactions suggest that backbone contacts dominate the interactions between TMEM55B and its binding partner. Mutation of G396^RL1^➔Leu abolishes the interaction with TMEM55B (**Figure 1F**). In addition to backbone interactions, two key residues of RILPL1 mediate sequence-specific interactions with TMEM55B. The first is E394^RL1^, whose sidechain makes a hydrogen bond with the backbone of K141^TM^. The second is E398^RL1^, which forms an electrostatic interaction with R151^TM^ (**Figure 1B-E**).

Mutagenesis of E398^RL1^➔Lys abolishes complex formation in cell-based co-immunoprecipitation (co-IP) assays (**Figure 1F**). Our understanding of these two glutamates is that they provide ‘anchors’ that mediate sequence-specific recognition of TMEM55B. There are no significant conformational changes upon RILPL1 binding, suggesting that the two RING domains are a rigid platform for protein interactions. The quality of refined electron density maps reveal confidence in RILPL1 peptide model building (**Suppl Figure S1**). Statistics of the data and refinement for the complexed RILPL1/TMEM55B and uncomplexed TMEM55B are shown in **Table 1**.

**Table 1:**
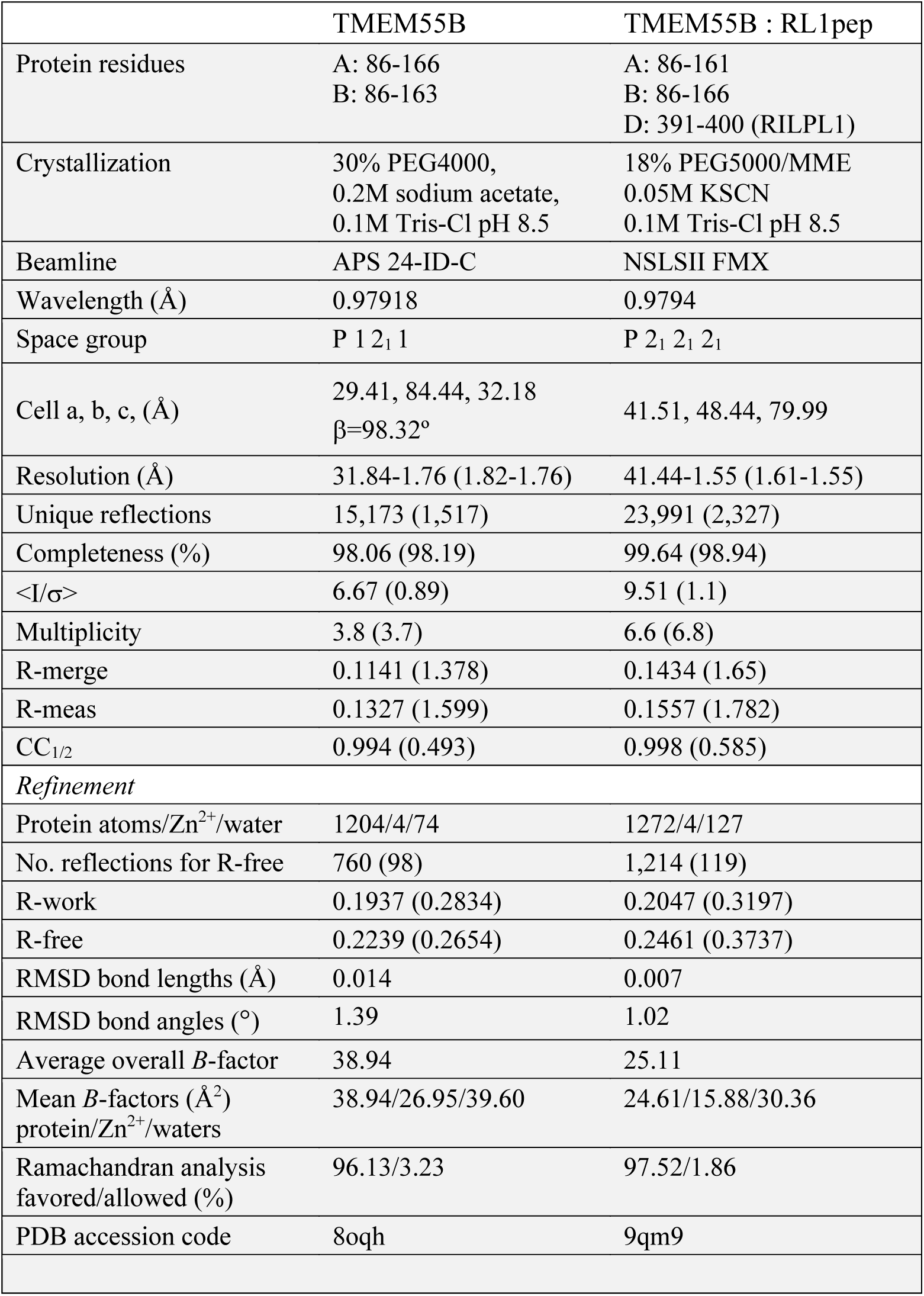
Data collection and refinement statistics

Although the overall complex is dominated by polar interactions that are largely sequence-independent, the C-terminal part of RILPL1 forms a non-polar contact that is also critical for complex formation. The sidechain of L400^RL1^ fits into a shallow pocket formed by the interface between the two RING-like domains. Mutagenesis of L400^RL1^➔Ala abolishes the interaction in co-IP assays^9^. The stoichiometry of the complex TMEM55B:RILPL1 peptide is 1:1, as evidenced by size exclusion chromatography coupled to multi-angle light scattering (SEC-MALS; **Suppl Figure S2**). Thermodynamic analyses of the interaction by isothermal titration calorimetry (ITC) reveals an enthalpically favourable interaction (**Δ**H= -12 kcal/mol) with a K_d_ of 700 nM (**Figure 1G**). Since RILPL1 is a dimer of parallel coiled coils through its RH1 and RH2 domains^8,14,15^, two TMEM55B molecules could in principle bind to a dimer of RILPL1.

### Comparisons of TMEM55B with the RBR E3 ligase RNF216

Comparisons of the structure of TMEM55B using FoldSEEK^16^ and DALI^17^ reveal similarities to RING domains of E3 ligases. The highest hit from FoldSEEK is RNF216 (**Figure 2A**), a Ring-between-Ring (RBR) E3 ligase [PDB code 7m4m^18^]. The root-mean-square (RMS) deviation of the superposition using only the C_α_ atoms of the C-terminal RING-like domain of TMEM55B is 1.2Å over 35 residues. Following a structure-based sequence alignment using Chimera^19^, there are 26% identities to RNF216 (residues 600-634; **Figure 2B**). The second highest structural hit is LysW from *Thermus thermophilus* [PDB code 5eio^20^], an amino group-carrier protein associated with lysine biosynthesis. There are no significant sequence identities with LysW apart from conserved Cys residues in TMEM55B. Interestingly, the E3 ligase Parkin is also identified by FoldSEEK as having structural similarities to TMEM55B. Although TMEM55B appears to have a structurally conserved RING-like fold, it appears to be unique among known proteins in having two tandem domains as a single globular unit that serve to facilitate the recognition of a RILPL1 peptide motif.

**Figure 2:**
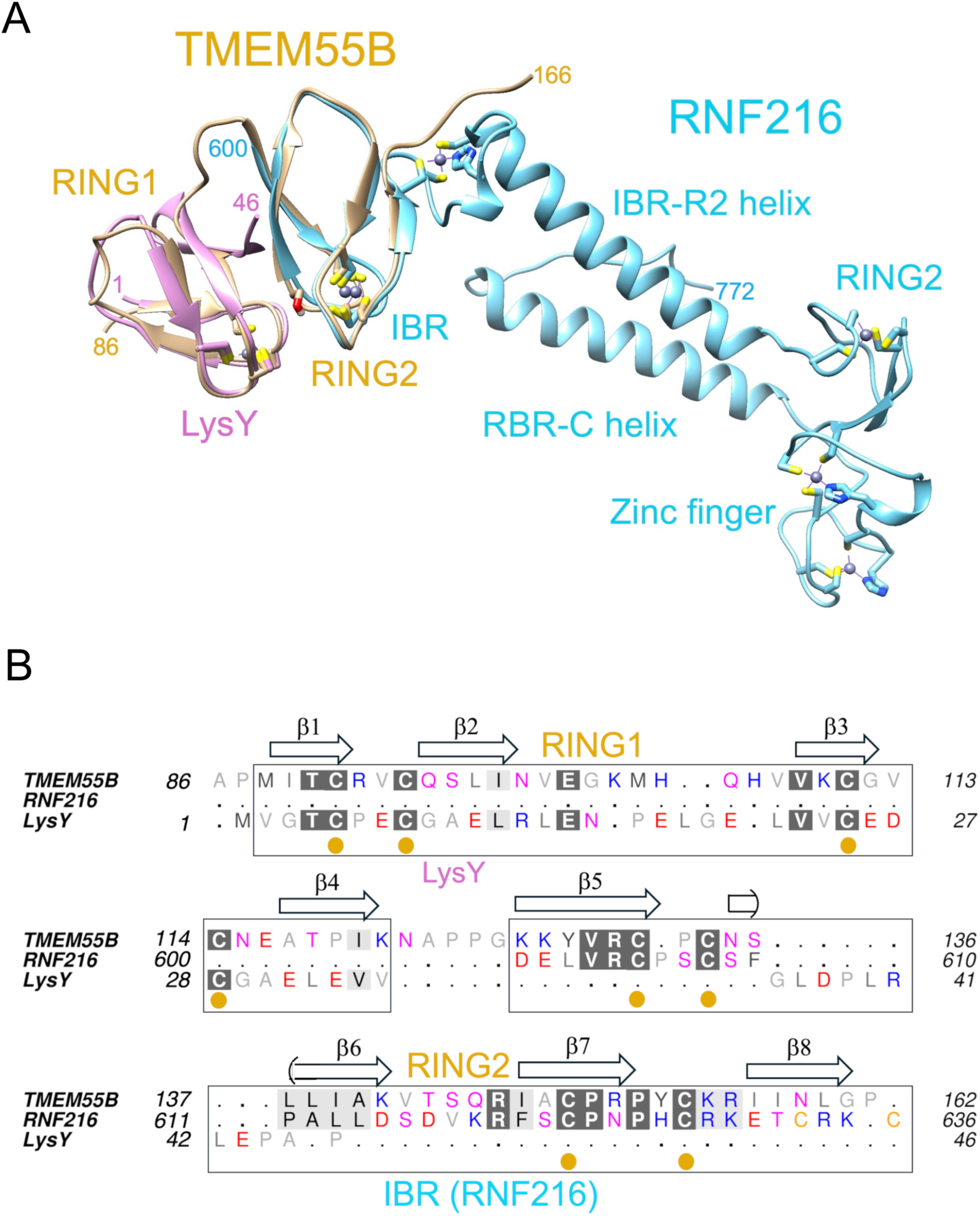
Comparisons of TMEM55B with known structures using FoldSEEK. **(A)** Least-squares superposition of RNF216 (blue) and LysW (magenta) with TMEM55B (yellow), shown as ribbon models. The side chains of cysteine residues that coordinate Zn2+ atoms within the RING domains and zinc finger are sticks. **(B)** Structure-based sequence comparisons of the proteins. The secondary structure annotations above the alignment corresponds to TMEM55B.

### Identification of new interactors of TMEM55B

The obligatory interaction between R151^TM^ and RILPL1 was exploited to find additional proteins that can recognize TMEM55B. DIA-MS analysis using the immunoprecipitate of TMEM55B WT and R151E^TM^ in the presence of LRRK2 Y1699C (kinase active mutant) and Rab8A Q67L (GTP-bound mutant) identified a set of interacting proteins which use the same (or overlapping) interface of TMEM55B (**Figure 3A**). In addition to RILPL1, a subset of strong interactors included JIP3, JIP4, OCRL, and WDR81 (**Figure 3B**), as evidenced by a clear contrast between WT TMEM55B and the R151E^TM^ mutation. To investigate further whether these interactions require phosphorylated Rab8A, we performed an additional mass spectrometry experiment transfecting Halo-TMEM55B WT with Rab8A Q67L either in the presence of LRRK2 Y1699C (kinase active mutant) or, LRRK2 Y1699C with D2017A (kinase dead mutant; **Figure 3C**). The Y1699C LRRK2 enhances the levels of pRab8A, while the inactive D2017A variant is expected to have low levels of pRab8A. TMEM55B enrichment in the presence of pRab8A is only maintained with RILPL1, not the other interactors (**Figure 3C**). Therefore, the binding of TMEM55B to JIP3, JIP4, WDR81 and OCRL is independent of phosphorylated Rab8. In contrast RILPL1 binding to TMEM55B is increased upon hyperphosphorylation of Rab8A (**Figure 3D**). The dependency of the cellular RILPL1 interaction with TMEM55B on phosphorylated Rab8A was further validated by co-IP and western blot analysis (**Figure 3E**). Overexpression of kinase dead D2017A/Y1699C double mutant of LRRK2 significantly reduced pRab8A levels and concomitantly led to >50% reduction in RILPL1 binding to TMEM55B (left panel, **Figure 3E**). Interestingly, Rab12 also exhibited increased co-IP with Halo-TMEM55B in a manner that was dependent upon LRRK2 activity (**Figure 3B**) and inhibited by the R151E^TM^ mutation in TMEM55B (**Figure 3A**). While the basis for Rab12 binding, is unclear, it is unlikely to reflect direct binding between Rab12 and TMEM55B. We speculate that Rab12 may share a common interactor with pRab8A, RILPL1 or TMEM55B, but this requires further investigation.

**Figure 3:**
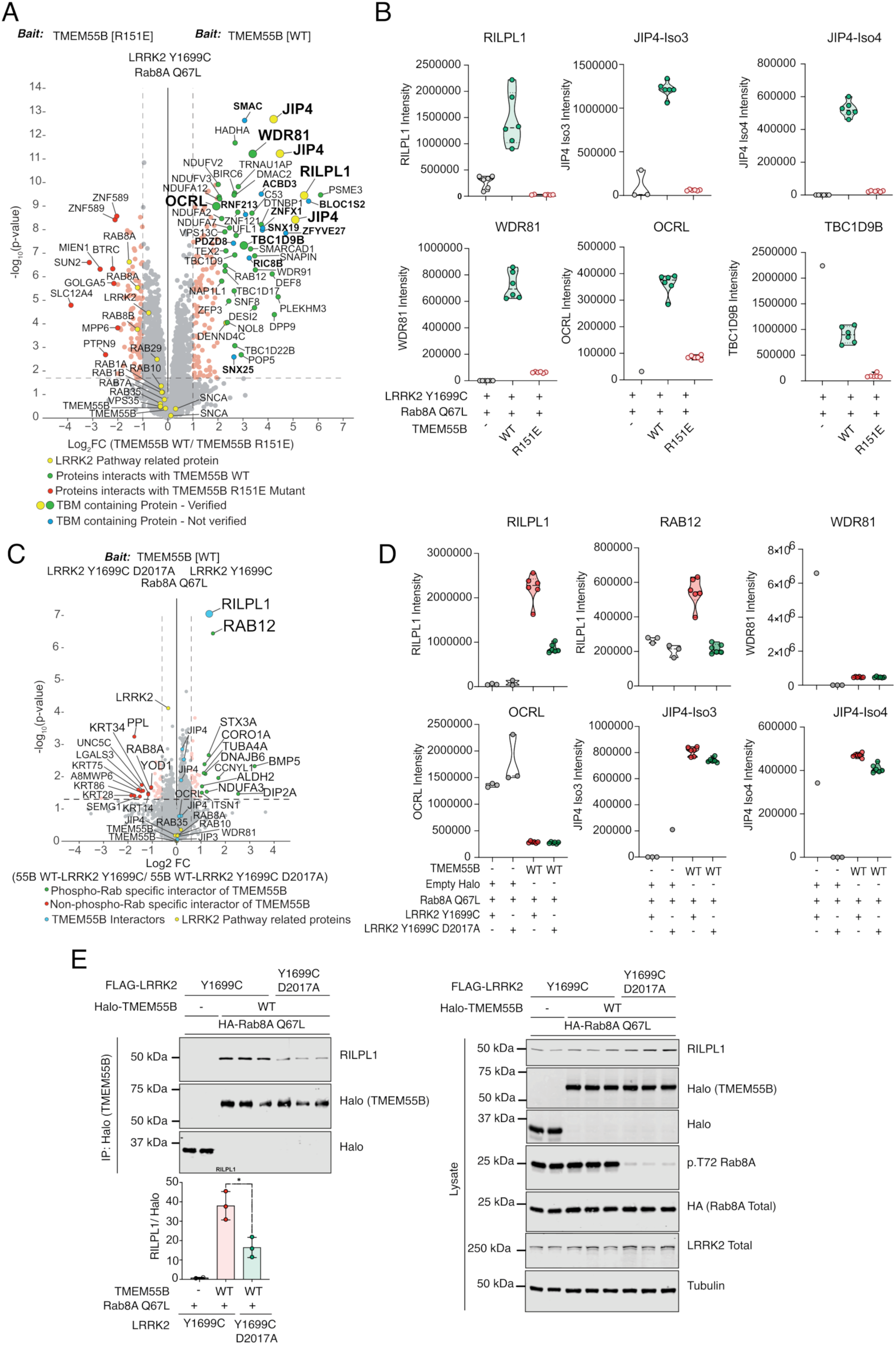
**(A)** Volcano plot depicting the fold-enrichment of proteins between IPs from Halo-TMEM55B WT and Halo-TMEM55B R151E mutant in the presence of LRRK2 Y1699C and Rab8A Q67L (two-tailed Student’s *t*-test, which is adjusted by permutation-based FDR of 5%). Curtain link, https://curtain.proteo.info/#/ed948504-461d-4e7b-b883-85268a8fb745. **(B)** Violin plots of the levels of indicated proteins. **(C)** Volcano plot depicting the fold-change of proteins between IPs from Halo-TMEM55B WT in the presence of either LRRK2 Y1699C (kinase active mutant) or LRRK2 Y1699C D2017A (kinase dead mutant) and Rab8A Q67L (*p*-value adjusted by two-tailed Student’s *t*-test, which is corrected by permutation-based FDR of 5%). Curtain link, https://curtain.proteo.info/#/b6bdc307-cfcb-418e-a232-1b9adde7a3ac. **(D)** Violin plots of the levels of RILPL1 and selected interactors of TMEM55B from (A). **(E)** HEK293 TMEM55B/A double knock-out cells were transiently transfected with the indicated proteins and lysed 24h post transfection. Halo-TMEM55B immunoprecipitations (right panel) or cell extracts (left panel) were subjected to quantitative immunoblot analysis using the LI-COR Odyssey CLx Western Blot imaging system and indicated antibodies. Quantitation of immunoblotting data (performed using ImageStudioLite software version 5.2.5, RRID:SCR_013715).

### TMEM55B Binding Motif is widespread among proteins linked to membrane trafficking

Our next goal was to investigate whether the TMEM55B Binding Motif (TBM), that we previously identified in RILPL1, is present in these new interactors of TMEM55B. The sequences of JIP3, JIP4, OCRL, and WDR81 were manually searched for the Ex(G/A)xEA motif, where ‘x’ represents variable residues. Further considerations were the following: (i) some degree of evolutionary conservation, and (ii) region of the polypeptide should be relatively flexible and surface-exposed. Motifs that were predicted to form inaccessible parts of α-helices or β-sheets were excluded as candidates. Possible TBM sequences from JIP3, JIP4, OCRL, WDR81, and TBC1D9B are aligned with RILPL1 (**Figure 4A**). Based on this alignment, we generated double mutants to abolish the interactions between these proteins and TMEM55B. The critical glutamates (equivalent to the E398^RL1^) were mutated to lysine, and A399^RL1^ equivalent residues were changed to leucine for JIP3, JIP4, OCRL, WDR81, TBC1D9B. Co-IPs from transient transfections in HEK293 cells revealed substantially reduced interaction of TMEM55B with mutants of JIP3, JIP4, OCRL, WDR81, and TBC1D9B (**Figures 4B-F**).

**Figure 4:**
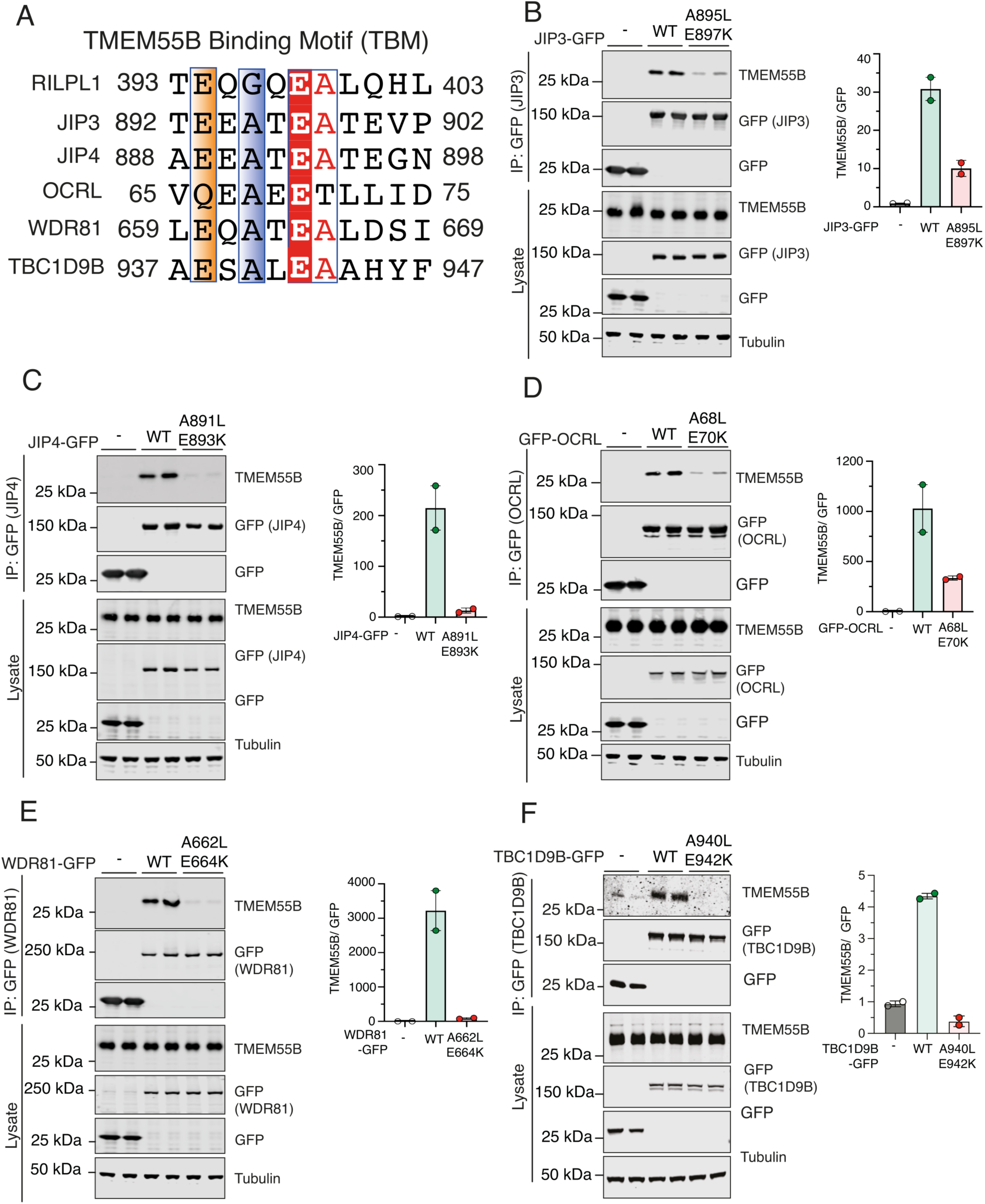
**(A)** Multiple sequence alignment of the TBMs of RILPL1, JIP3, JIP4, OCRL, WDR81, and TBC1D9B using Muscle5 (http://www.drive5.com/muscle/) and ESPript. Conserved Glycine/Alanine residue and Glutamic acid residue were highlighted in blue, and red. **(B to F)** HEK293 wild type cells were transiently transfected with the indicated proteins and lysed 24h post transfection. GFP immunoprecipitations (upper panel) or cell extracts (lower panel) were subjected to quantitative immunoblot analysis using the LI-COR Odyssey CLx Western Blot imaging system and indicated antibodies. Quantitation of immunoblotting data (performed using ImageStudioLite software version 5.2.5, RRID:SCR_013715). Quantitation of immuno-blotting data (performed using ImageStudio Lite software version5.2.5, RRID:SCR_013715) is shown as mean ± SEM.

### *In vitro* pulldowns of TMEM55B by TBM peptides

From a structural perspective, the interactions of TMEM55B with JIP3, JIP4, OCRL, and WDR81 appear to involve internal TBM sequences within these proteins, rather than the C-terminal TBM employed by RILPL1 (**Figure 5A**). Among these, JIP4 is particularly noteworthy. Structural predictions using AlphaFold3 suggest that JIP4 may form additional contacts with TMEM55B beyond the core TBM region (**Suppl Figure S3**). To probe this further, we mutated Lys103 to Glu (K103E) within TMEM55B. This mutation modestly enhanced RILPL1 binding but dramatically impaired JIP4 interaction, supporting the notion that JIP4 engages TMEM55B through additional structural features beyond the canonical TBM (**Suppl Figure S3**). In addition to these validated interactors, several other proteins, including BLOC1S2, ZFYVE27, ACBD3, RIC8B, SNX19, SNX25, PDZD8, RNF213, ZNFX1, and SMAC were also enriched in immunoprecipitates of wild-type TMEM55B, but not with the TBM-binding-deficient R151E mutant. These are highlighted with sky blue circle in **Figure 3A**. Although these candidates were not investigated further in this study, sequence analysis indicates they may also contain potential TBMs (**Suppl Fig S4A**), suggesting they could represent additional TMEM55B interactors, but this would need to be experimentally validated in future work. The domain organization of TBC1D9B is also shown, with the TBM situated in a flexible region toward the C-terminus (**Suppl Fig S4B**).

**Figure 5:**
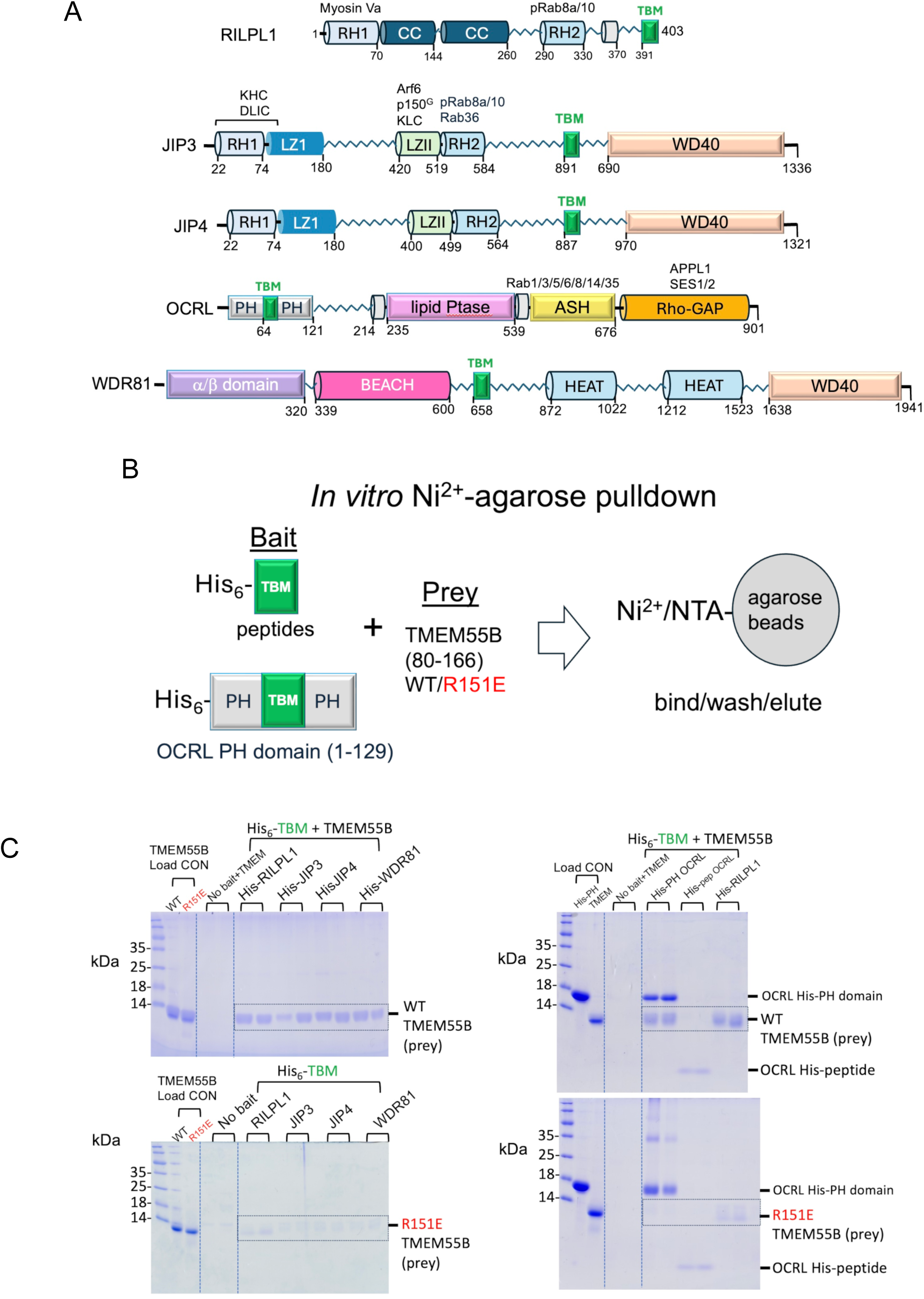
*In vitro* pulldowns of TMEM55B with TBM peptides. (A) Domain organization of TMEM55B interactors depicting the sequence context of the TBM. Regions that are α-helical have been drawn as cylinders; α/β folds are shown as rectangles; and unstructured segments are zigzag lines. (B) Cartoon depiction of the in vitro pulldown strategy using His-tagged peptides. **(C)** His-tagged peptides were used as bait to successfully pulldown untagged WT TMEM55B region 80-166, which migrates at approximately 10kDa (top panel). As a control, the R151E variant of TMEM55B was used, which abolishes the interactions with RILPL1 (bottom panel). Similarly, the His-tagged PH domain of OCRL1 (1-129, ∼16kDa) was used as bait to successfully pulldown WT TMEM55B (top) or the R151E variant (bottom). As a further control, a peptide corresponding to the TBM of OCRL failed to pulldown TMEM55B (top panel). Protein/peptide concentrations were 10 μM (see Methods) and visualization was enabled by Coomassie Brilliant Blue G250.

Given that the TBMs reside in flexible regions of the adaptor proteins (**Figure 5A**), peptides should be able to mimic their interactions with TMEM55B. Peptides corresponding to the potential TBMs were synthesized with an N-terminal hexahistidine tag to enable *in vitro* pulldowns of TMEM55B (80-166; **Figure 5B**). Slightly extended peptides beyond the minimal TBMs were generated to preclude any structural or conformational restrictions on the interaction with TMEM55B. The peptide TBM from RILPL1 was used as a positive control, and the mutant R151E^TM^ of TMEM55B provided a negative control. Peptides run on conventional SDS-PAGE gels are generally not visible in this assay.

Pulldowns using His-tagged RILPL1 peptide showed a clear band for untagged TMEM55B (**Figure 5C**). The pulldown was abolished when the R151E^TM^ variant of TMEM55B was used. RILPL1 provides a qualitative assessment of pulldown interactions under these conditions, since it has a known K_d_ of ∼700nM. Pulldowns with peptides from other interacting partners revealed that JIP3, JIP4, and WDR81 also interact with TMEM55B, showing clear contrast between WT and R151E^TM^ variants of TMEM55B (**Figure 5C**). OCRL is an interesting case – the His-tagged peptide did not significantly interact with TMEM55B, relative to RILPL1 (**Figure 5C**, *right panel*). The predicted TBM is situated in a loop region flanked by two anti-parallel β-strands (63-82) in the N-terminal PH domain of OCRL. To further explore this interaction, we performed a pulldown with a His-tagged variant of the PH domain (residues 1-129 of OCRL). Data revealed that, in contrast to the OCRL peptide alone, the PH domain interacts robustly with TMEM55B (**Figure 5C**). OCRL is also distinct in having a glutamine (Q66) as the first anchor of the TBM, instead of glutamate (**Figure 4A**). The side chain is identical in length and the amide would, in principle, be able form a hydrogen bond with the backbone of K141^TM^. It is possible that, like JIP4, additional interactions beyond the TBM are associated with complex formation.

### Isothermal titration calorimetry with TBM peptides and OCRL PH domain

The affinities between TBMs and TMEM55B were measured using isothermal titration calorimetry (**Figure 6**). Collectively, the K_d_ values for TBMs from RILPL1, OCRL, WDR81, JIP3 and JIP4 ranged from 20nM to 2900 nM, and all titrations revealed a favorable enthalpy (**Δ**H from -6 to -12 kcal/mol; **Figure 6** and **Table 2**). The errors in parameters were calculated from fitting of the curves shown using Origin software, as described (see Methods). All of the individual experiments and associated fitting statistics are in Table 2. Overall, these data demonstrate that despite sequence diversity the TBMs are capable of interacting with TMEM55B. However, the specific thermodynamic parameters are unlikely to reflect affinities of complexes in a cellular context. While the key salt bridge and backbone hydrogen bonds are likely conserved, differences in the surrounding architecture among these trafficking adaptors and their relative abundance at the lysosome will influence the strength of interactions.

**Figure 6:**
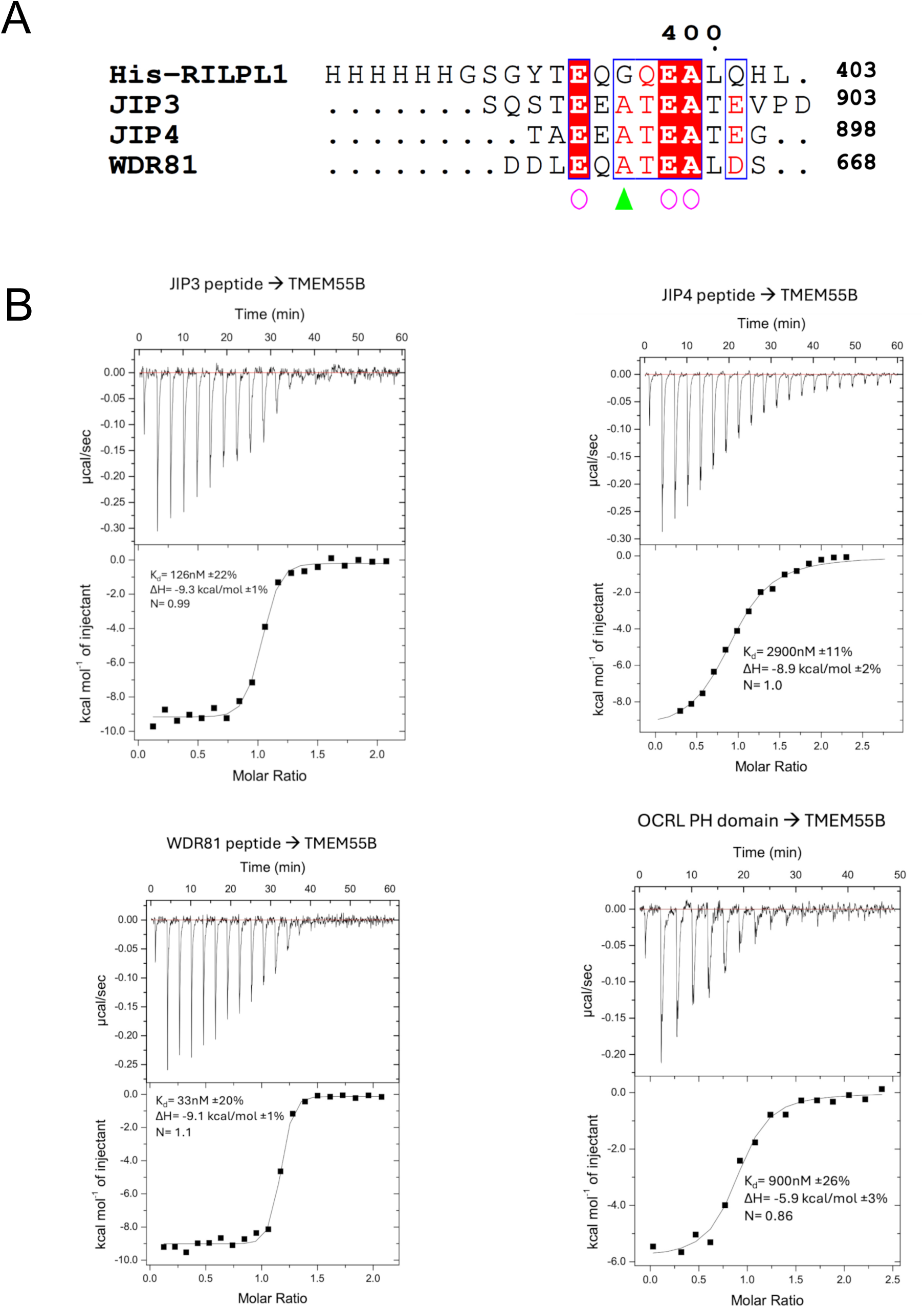
Thermodynamics of the interactions between TBMs and TMEM55B using isothermal titration calorimetry. **(A)** Sequences of the peptides used for the experiment are aligned. **(B)** Peptides corresponding to the TBMs of JIP3, JIP4 and WDR81 were injected into a cell containing TMEM55B (residues 80-160). For OCRL, the PH domain was purified as described in METHODS and used for injections into the cell.

**Table 2:** Thermodynamics of TMEM55B binding to interactors

|  | **K<sub>d</sub> (nM)** | **ΔH (kcal/mol)** | **ΔS (cal/K•mol)** | **N** |
| --- | --- | --- | --- | --- |
| **RILPL1 391-403** |  |  |  |  |
| (i) 0.45mM→ 0.06mM | 590 ±7% | -12.1 ±5% | -12.7 | 0.95 |
| (ii) 0.3 mM→0.04mM | 1000 ±22% | -13.6 ±22% | -19 | 0.87 |
| (iii) 0.45mM→0.045mM | 680 ±23% | -11.4 ±5% | -10.7 | 0.87 |
| **OCRL-PH domain** |  |  |  |  |
| (i) 0.3mM→0.015mM | 1700 ±50% | -5.3 ±10% | 8.6 | 0.69 |
| (ii) 0.35mM→0.03mM | 900 ±26% | -5.9 ±3% | 7.6 | 0.86 |
| (iii) 0.5mM→0.05mM | 460 ±100% | -6.7 ±10% | 6 | 0.68 |
| **WDR81 (657-668)** |  |  |  |  |
| (i) 0.3mM→0.04mM | 20 ±27% | -9.0 ±1% | 4.4 | 0.94 |
| (ii) 0.2mM→0.02mM | 33 ±20% | -9.1 ±1% | 3.2 | 1.1 |
| (iii) 0.2mM→0.02mM | 12 ±92% | -8.7 ±2% | 6.4 | 1.1 |
| **JIP3 (889-903)** |  |  |  |  |
| (i) 0.3mM→0.03mM | 127 ±47% | -8.2 ±3% | 3.4 | 1.2 |
| (ii) 0.3mM→0.03mM | 126 ±22% | -9.3 ±1% | -0.2 | 0.99 |
| (iii) 0.3mM→0.03mM | 103 ±35% | -9.3 ±2% | 0.2 | 1.1 |
| **JIP4 (887-897)** |  |  |  |  |
| (i) 0.4mM→0.03mM | 2900 ±11% | -8.9 ±2% | -5.1 | 1.0 |
| (ii) 0.4mM→0.03mM | 2200 ±13% | -9.7 ±8% | -7.1 | 0.91 |
| (iii) 0.4mM→0.03mM | 3100 ±37% | -8.7 ±10% | -4.5 | 0.78 |

The exquisite specificity of the TMEM55B surface toward RILPL1 was demonstrated by a series of single-site mutations to alanine (**Suppl Figure S5**). Mutant E116A^TM^ abolished the interaction – this glutamate forms a salt bridge with R151^TM^ and presumably helps orient the guanidino side chain for a salt bridge with E398^RL1^ (**Figure 1**). Similarly, L137A^TM^ abolished the interaction with RILPL1. The side chain of L137^TM^ is within van der Waals distance of the backbone of G396^RL1^.

## DISCUSSION

Here we report the structural characterization of a motif, ‘E-x-G/A-x-E-A’, that enables binding to lysosome-localized TMEM55A/B. The interactions are driven by backbone hydrogen bonds and two glutamates with an intervening Gly/Ala small residue. The TBM forms a terminal β-strand and bridges the two tandem RING-like domains across a relatively shallow surface of TMEM55B. A further requirement for this TBM is its presence in a flexible stretch of polypeptide, free from backbone constraints, to facilitate intimate backbone hydrogen bonds with TMEM55B. Thermodynamics, structural and cellular assays are consistent with a model in which LRRK2 phosphorylation of Rab8A enables RILPL1 recruitment to TMEM55B-associated lysosomes. The C-terminal RILPL1 TBM is in a flexible region far from the RH2 domain which recognizes phospho-Rab8A/10 (**Figure 7**). Therefore, it is unlikely that phospho-Rab binding can directly influence RILPL1/TMEM55B interactions *via* allostery or an overlapping 3-way interface.

**Figure 7:**
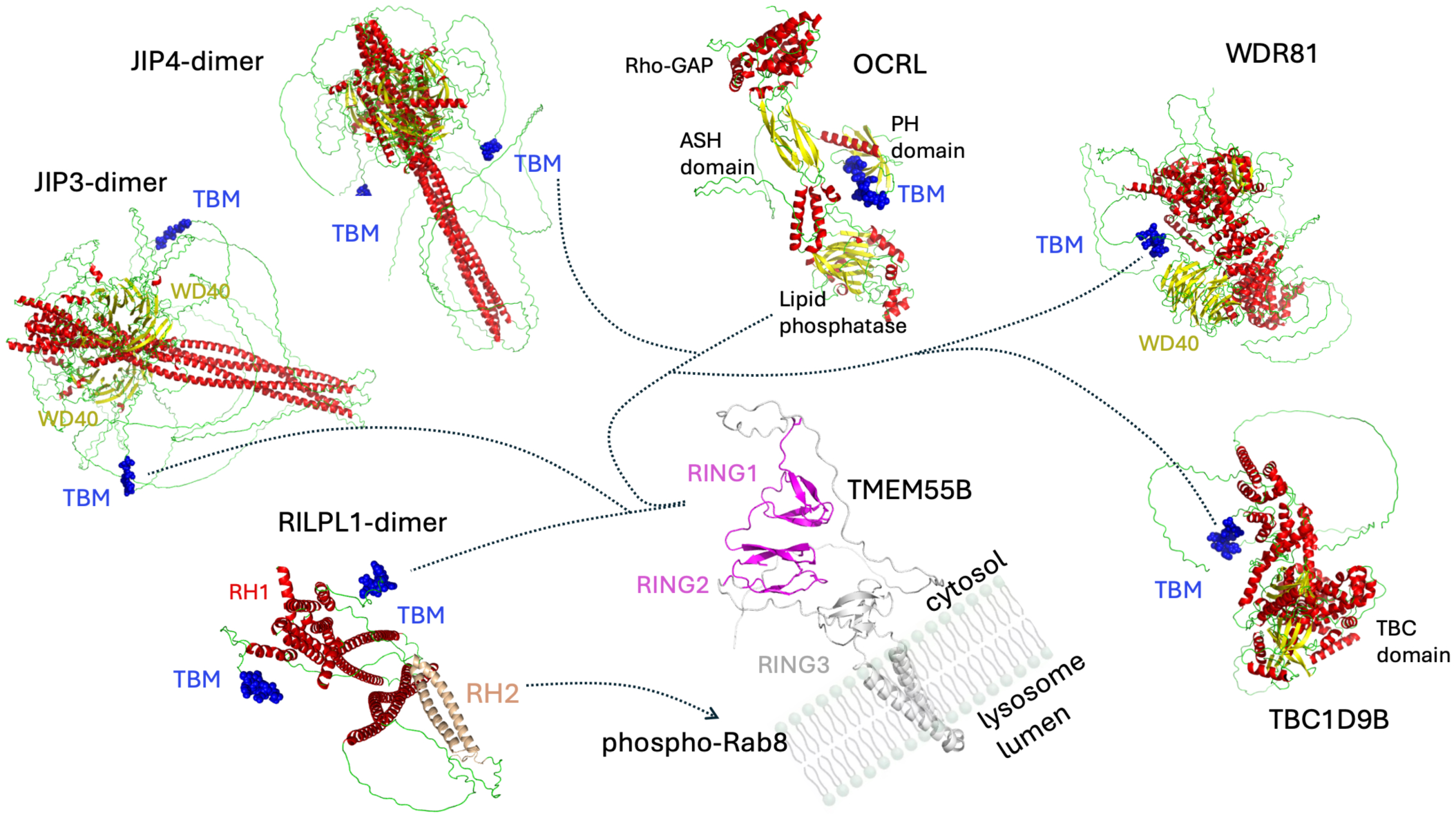
Model for TBM mediated interactions with TMEM55B. RILPL1 complex formation is dependent on recruitment to lysosomes by phosphorylated Rab GTPases *via* its RH2 domain (peach colour). However, interactions with TBMs in other proteins are independent of LRRK2. TBMs are shown as blue van der Waals spheres, and colour annotation follows secondary structure, with α-helices in red, β-sheets in yellow, and unstructured regions in green. Models were predicted using the AlphaFold3 server and figures generated with PyMol^30^.

We have also identified the TBMs of JIP3, JIP4, WDR81, OCRL, TBC1D9B and possibly in other proteins (**Figure 7** and **Suppl Figure S4**). Like RILPL1, JIP3 and JIP4 also have an RH2 domain that binds to phospho-Rab10^8,14^. However, only RILPL1 complexes with TMEM55B are dependent on phosphorylated Rab GTPases. The peptide motifs reside in highly flexible and exposed segments between globular domains. The common feature of TMEM55B interactors is their link to various aspects of membrane trafficking. JIP3/4 are adapters for microtubule motors for transport of cargo including vesicles, protein, mRNA and they also regulate organelle dynamics ^21,22^. In JIP3, the TBM is situated within a disordered region between the N-terminal α-helical coiled coils and a predicted propeller-like C-terminal WD40 domain. Intriguingly, RILPL1 and JIP4 regulate opposing processes that involve formation/breakdown of tube-like vesicles from damaged lysosomes^23^. WDR81 is a 1,941-residue protein which also has a WD40 domain (residues 1,638-1,941) at its C-terminus. There are no experimental structures of the protein, but the AlphaFold3 model suggests the presence of multiple α-helical domains flanking the TBM (residues 658-668). WDR81 functions are associated with autophagy, endosomal trafficking, and its loss of function causes severe microcephaly in patients^24,25^.

The precise function of TMEM55B remains elusive despite a structural relationship to the RING-like domains of RBR E3 ligases. It is unlikely to be a conventional E3 ligase for the following reasons: (i) RING1 and RING2 domains have no intervening regions and act as a tandem single domain, which is not typical of substrate binding regions, and (ii) there is no catalytic cysteine in the predicted membrane-proximal RING3 domain (**Figure 7**). However, an atypical membrane-bound E3 ligase cannot formally be excluded as the biological function of TMEM55B. Complicating the functional role of TMEM55B, the native full-length protein may be oligomeric. While AlphaFold3 does not provide a clear indication of oligomerization, future studies could explore whether TMEM55B is capable of assembling into oligomeric complexes. Therefore, TMEM55B may provide a signaling hub at the lysosome to bridge multiple cytoskeletal networks. The structural studies presented here, together with 3-D predictions, also suggest that OCRL and JIP4 complexes may involve additional motifs beyond the TBM. In the future, it may be possible to tease out specific roles of TMEM55B interactors through mutagenesis and cellular assays. Taken together, these findings indicate that TMEM55B functions as a central hub for adaptor protein recruitment at the lysosomal membrane. Our study uncovers previously unrecognized regulatory networks linked to TMEM55B and lysosomal function, highlighting new avenues for future investigation.

## ACKNOWLEDGMENTS

We thank Dr Thomas Macartney for designing the CRISPR-Cas9 related constructs and the excellent technical support of the MRC-PPU DNA sequencing service (coordinated by Gary Hunter), the MRC-PPU tissue culture team (coordinated by Dr Edwin Allen), the MRC-PPU mass spectrometry facility team (coordinated by Dr Renata Soares) and the MRC-PPU Reagents and Services team (coordinated by Dr James Hastie). A.R.K. was funded by Research Ireland through a Research Frontiers grant, 20/FFP-A/8446. DRA was supported by the UK Medical Research Council. This study was funded by the joint efforts of The Michael J. Fox Foundation for Parkinson’s Research (MJFF) and Aligning Science Across Parkinson’s (ASAP) initiative. MJFF administers the grant (ASAP-000463) on behalf of ASAP and itself to D.R.A. This research used resources of the Advanced Photon Source, a U.S. Department of Energy (DOE) Office of Science user facility operated for the DOE Office of Science by Argonne National Laboratory under Contract No. DE-AC02-06CH11357. This work is based in part upon research conducted at the NE-CAT beamlines at the Advanced Photon Source, which are funded by the National Institute of General Medical Sciences from the National Institutes of Health (P30 GM124165). This research used beamline FMX of the National Synchrotron Light Source II, a U.S. Department of Energy (DOE) Office of Science User Facility operated for the DOE Office of Science by Brookhaven National Laboratory under Contract No./ DE-SC0012704. The Center for BioMolecular Structure (CBMS) is primarily supported by the National Institutes of Health, National Institute of General Medical Sciences (NIGMS) through a Center Core P30 Grant (P30GM133893), and by the DOE Office of Biological and Environmental Research (KP1605010).

## Data and materials availability

All primary data presented here have been deposited in publicly accessible repositories. Immunoblotting data have been deposited in Zenodo, namely, main figure and supplementary immunoblotting images (https://doi.org/10.5281/zenodo.16883734), and GraphPad files for all figures (https://doi.org/10.5281/zenodo.16883557). Structural data has been deposited in the Protein Data Bank with accession codes 8oqh (TMEM55B) and 9qm9 (RILPL1/TMEM55B complex). SEC-MALS data (https://doi.org/10.5281/zenodo.16814518), protein gels from in vitro pulldowns (https://doi.org/10.5281/zenodo.16814483) and ITC data files compatible with Origin software have also been deposited (https://doi.org/10.5281/zenodo.16814276). Proteomic data have been deposited in the ProteomeXchange PRIDE repository (identifiers: PXD067414) and Zenodo (https://doi.org/10.5281/zenodo.16881244). All plasmids and antibodies generated at the MRC PPU at the University of Dundee can be requested through our website: https://mrcppureagents.dundee.ac.uk/. On this website, type in the name of the protein required in the search box, and the list of available plasmids, proteins, and antibodies available against that protein is displayed. Select those of interest and complete ordering form. Please note that the request of plasmid requires a universal material transfer agreement (MTA) that can be completed online at the time of plasmid request.

## AUTHOR CONTRIBUTIONS

A.R.K., P.P. and D.R.A. conceived, designed and A.R.K. and D.R.A. supervised the study. D.W. and A.R.K. performed all the protein purification, calorimetry, and crystallographic studies. P.P. performed all the cellular expression and co-IP experiments. P.P. performed and R.S.N. analysed the mass spectrometry related experiments; M.C. designed and generated the cDNAs related to co-IP experiments. J.S. performed the TBC1D9B related Co-IP validation experiment. A.R.K, P.P. and D.R.A wrote the manuscript with input from all co-authors. All authors read and approved this version.

## STAR METHODS

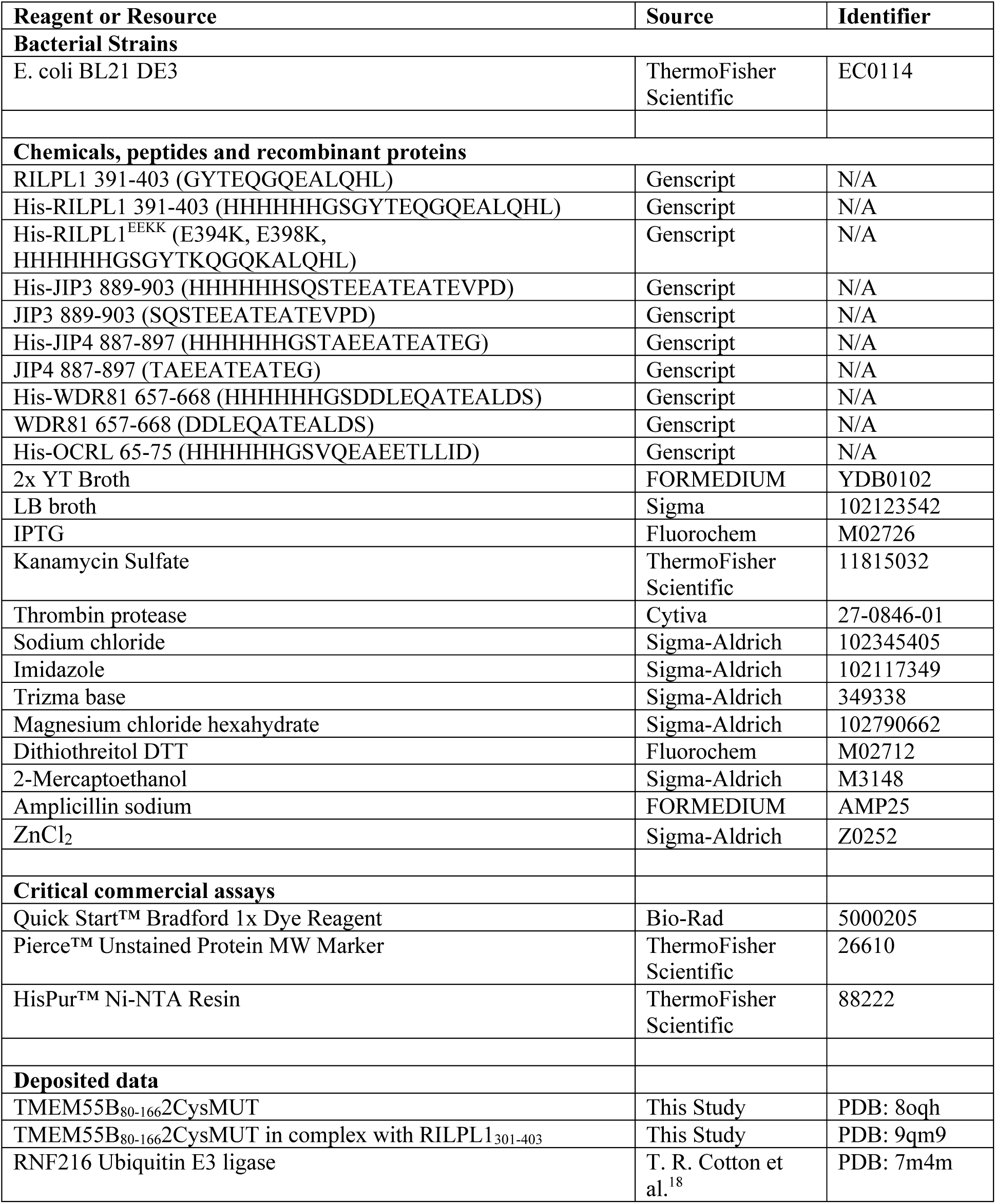
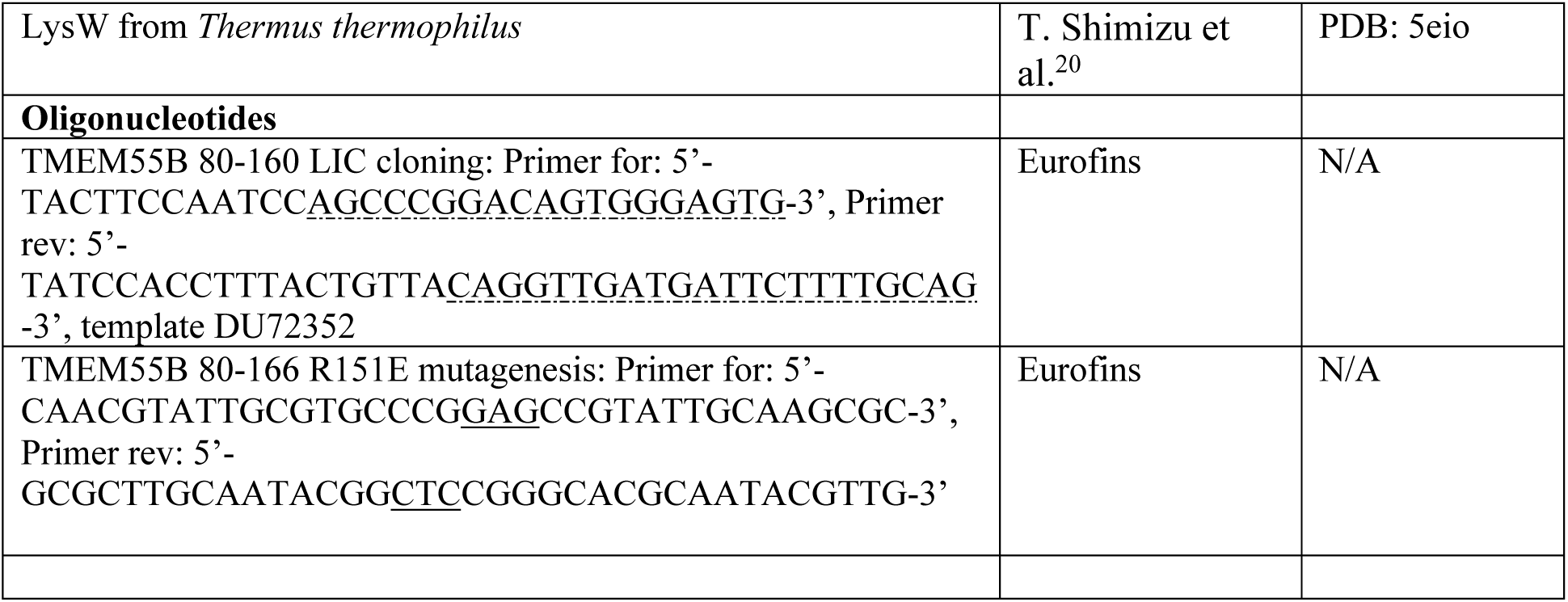
KEY RESOURCES TABLE

### Constructs used for cellular expression (https://mrcppureagents.dundee.ac.uk)

**Constructs used for cellular expression (https://mrcpppureagents.dundee.ac.uk)**
| **DU number:** | **Plasmid:** |
| --- | --- |
| DU13156 | pCDNA5 FRT/TO GFP Empty |
| DU72471 | pCDNA5 FRT/TO Halo Empty |
| DU26486 | pCMV5 FLAG-LRRK2 Y1699C |
| DU52703 | pCMV5 FLAG-LRRK2 Y1699C D2017A |
| DU51181 | pCDNA5 FRT/TO HA-Rab8A Q67L |
| DU27305 | pCDNA5 FRT/TO RILPL1-GFP |
| DU75853 | pCDNA5 FRT/TO RILPL1 G396L GFP |
| DU71241 | pCDNA5 FRT/TO RILPL1 E398K GFP |
| DU70318 | pCDNA5 FRT/TO Halo-TMEM55B |
| DU72569 | pCDNA5 FRT/TO Halo-TMEM55B R151E |
| DU72527 | pCDNA5 FRT/TO Halo-TMEM55B K110A |
| DU72528 | pCDNA5 FRT/TO Halo-TMEM55B E116A |
| DU72529 | pCDNA5 FRT/TO Halo-TMEM55B K127A |
| DU72530 | pCDNA5 FRT/TO Halo-TMEM55B R131A |
| DU72531 | pCDNA5 FRT/TO Halo-TMEM55B N135A |
| DU72532 | pCDNA5 FRT/TO Halo-TMEM55B L137A |
| DU72533 | pCDNA5 FRT/TO Halo-TMEM55B 3C L138A |
| DU72534 | pCDNA5 FRT/TO Halo-TMEM55B 3C I139A |
| DU72535 | pCDNA5 FRT/TO Halo-TMEM55B I147A |
| DU27721 | pCDNA5 FRT/TO JIP3-GFP |
| DU72982 | pCDNA5 FRT/TO JIP3 A895L E897K GFP |
| DU27684 | pCDNA5 FRT/TO JIP4-GFP |
| DU80198 | pCDNA5 FRT/TO JIP4 A891L E893K GFP |
| DU80129 | pCDNA5 FRT/TO GFP OCRL |
| DU80192 | pCDNA5 FRT/TO GFP OCRL A68L E70K |
| DU77478 | pCDNA5 FRT/TO WDR81-GFP |
| DU77495 | pCDNA5 FRT/TO WDR81 A662L E664K GFP |
| DU69593 | Sense A Guide RNA for TMEM55B CRISPR-Cas9 KO cell generation |
| DU69594 | Anti-sense A Guide RNA for TMEM55B CRISPR-Cas9 KO cell generation |
| DU69595 | G1 Single Guide RNA for TMEM55B CRISPR-Cas9 KO cell generation |
| DU69596 | G2 Single Guide RNA for TMEM55B CRISPR-Cas9 KO cell generation |
| DU69706 | Sense A Guide RNA for TMEM55A CRISPR-Cas9 KO cell generation |
| DU69708 | Anti-sense A Guide RNA for TMEM55A CRISPR-Cas9 KO cell generation |
| DU69710 | G1 Single Guide RNA for TMEM55A CRISPR-Cas9 KO cell generation |
| DU69711 | G2 Single Guide RNA for TMEM55A CRISPR-Cas9 KO cell generation |

### Peptides used in this study

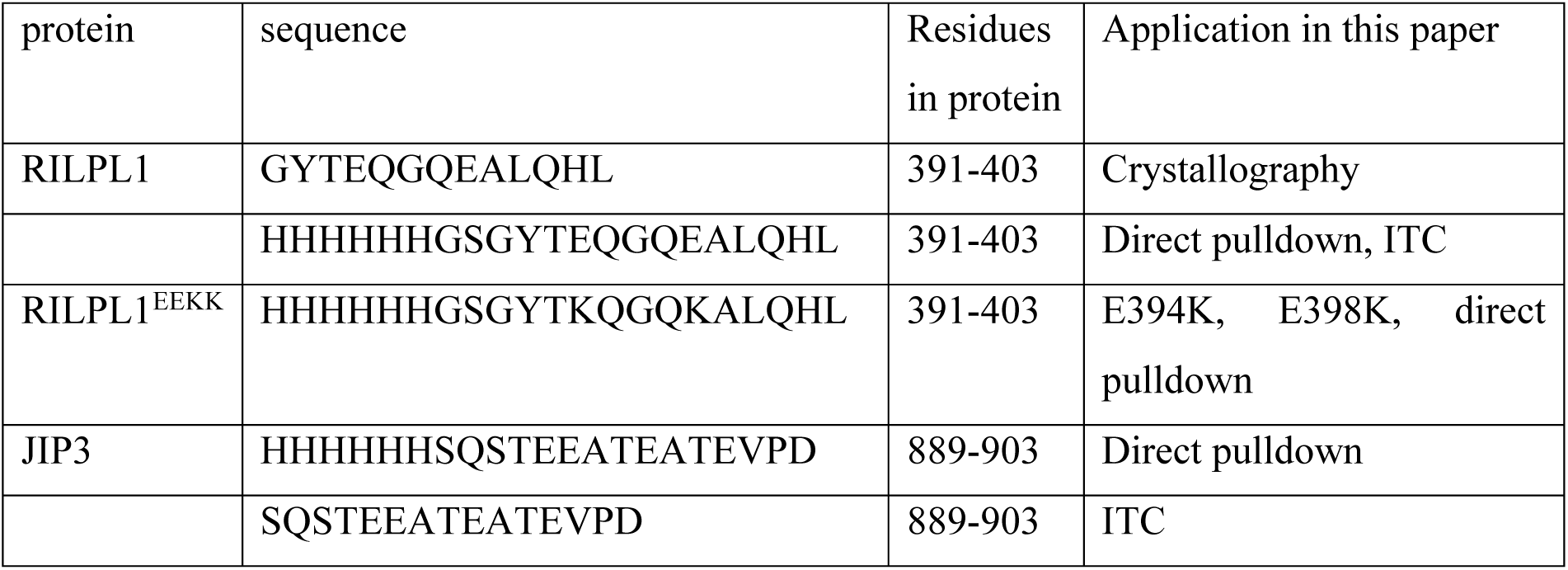
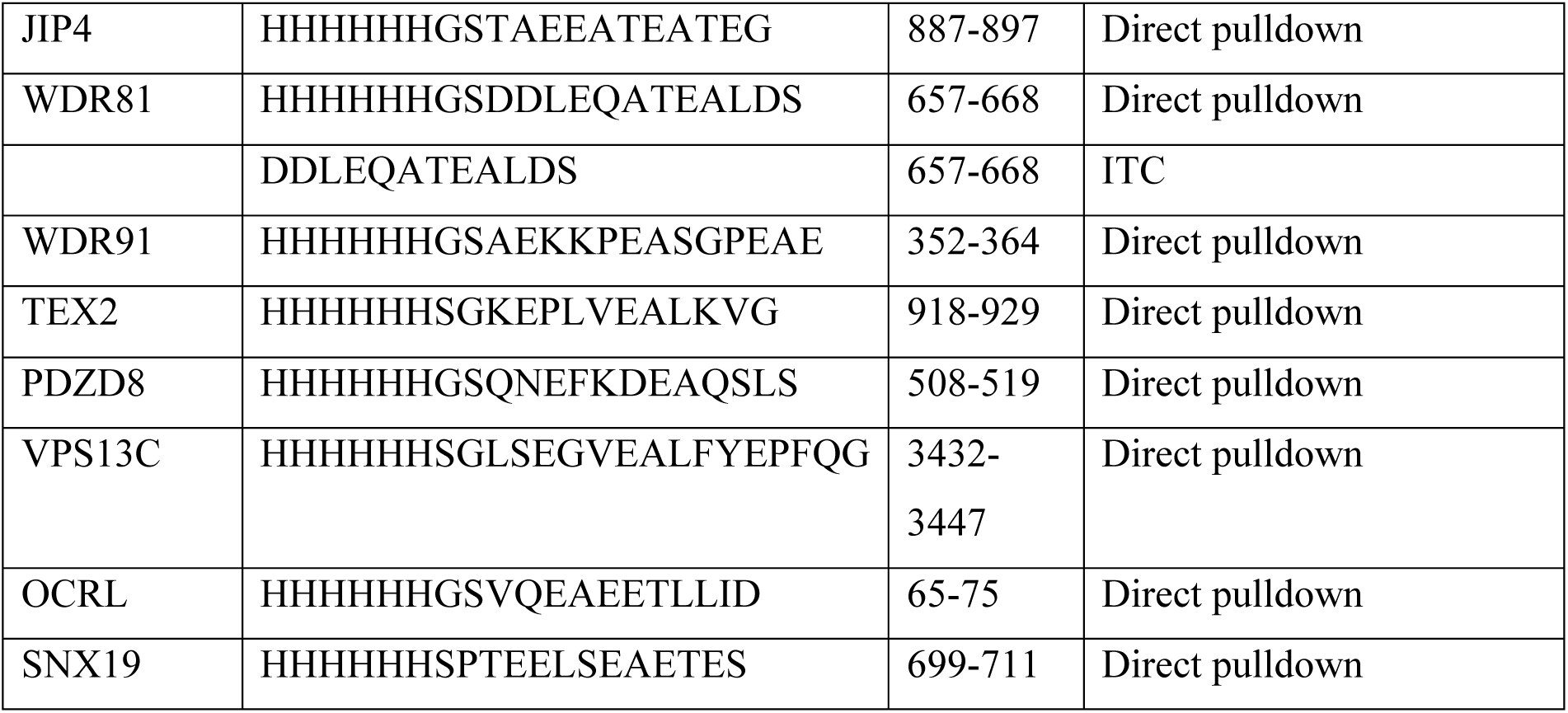

### Constructs for protein expression

TMEM55B constructs for recombinant proteins used for crystallography and static light scattering, isothermal titration calorimetry (ITC) or pulldown analyses were either (i) ordered from Genscript (Rijswijk, Netherlands) with the sequence optimized for expression in *E. coli*, (ii) subcloned using PCR amplification and subsequent ligation-independent cloning (LIC) cloning, or (iii) generated by site directed mutagenesis from an existing construct. All TMEM55B constructs and sequence numbering are based on UniProt entry Q86T03 isoform 2.

### Peptides for co-crystallization and ITC

All peptides used in this study were synthesized by Genscript (Rijswijk, Netherlands). Purity of the peptides for crystallography was >98%, others between 85-95%.

## METHOD DETAILS

### Recombinant expression and purification of TMEM55B

Plasmids encoding TMEM55B residues 80-166 were transformed into *E. coli* BL21(DE3) and maintained in LB medium. For expression, 500mL or 1L of 2xYT were inoculated at a 1:50 ratio with a saturated overnight culture and grown to an OD at 600nm of 0.6-0.8 at 37°C, 180rpm. The temperature was reduced to 18°C for ∼1h. The expression was induced by adding 500mM IPTG and supplementing 50µM ZnCl_2_. After overnight induction at 18°C cells were harvested by centrifugation. For purification of the recombinantly expressed protein extraction buffer (20mM Tris pH8.0 at RT, 300mM NaCl, 10mM Imidazole and 10mM β-mercaptoethanol) was added to the bacterial pellets. After resuspension and homogenization in a dounce homogenizer cells were disrupted by sonication. Insoluble material was separated from soluble proteins by 45min centrifugation, 20000 x g, 4°C. The supernatant was then added to a gravity flow column with a bed of Ni-NTA agarose (Thermo Fisher Scientific, Rockford, USA). The hexa-Histidine tagged TMEM55B construct bound to the resin was washed with extraction buffer followed by a more stringent wash (40mM Imidazole). The Elution occurred with 200mM Imidazole. Eluted proteins were dialyzed into extraction buffer. After 1h dialysis at 4°C 20 IU thrombin were added to the dialysis tube to cleave off the hexa-Histidine tag. Separation of cleaved from uncleaved protein was conducted by applying a second Ni-NTA agarose column. The final TMEM55B construct covered the residues 80-166 and include an N-terminal GSHM-peptide derived from the pET15b expression vector.

The production of crystallography-grade TMEM55B was finalized with a 50% low to high salt buffer (10mM Tris pH8.0 at RT, 10mM/1M NaCl, 1mM DTT) ion exchange chromatography using a MonoS 5/50GL column. The fractions containing the protein were subjected to gel filtration on a Superdex 75 10/300 column (buffer: 20mM Tris pH7.5 at RT/pH8.0 at 4°C, 150mM NaCl, 1mM DTT). The resulting fractions were analyzed by SDS-PAGE for purity, pooled and concentrated for crystallography. Proteins used in pulldown analyses, SLS and ITC were purified skipping the ion exchange chromatography step. The OCRL PH domain construct was expressed and purified using a similar protocol. However, no additional ZnCl_2_ was added to the expression media.

The above details of TMEM55B purification for structural and biophysical studies are also provided in dx.doi.org/10.17504/protocols.io.bp2l6yom1vqe/v1.

### Crystallization and X-ray data collection

TMEM55B_80-166_2CysMUT was concentrated prior to setting up in MD-2 sitting drop 96-well plates with various sparse-matrix crystallization screens. The resulting crystals were harvested in 30% glycerol directly from condition D1 of a “Structure Screen” (MD1-30, Molecular Dimensions). For the complex, proteins were concentrated to 20 mg/ml or higher. Screening conditions that revealed crystals were optimized and replicated in in 24 well plates as hanging drops. Best growth was obtained in the following condition: 0.05M KSCN, 0.1M Tris pH 8.5, 18% PEG5000/MME. Crystals were harvested in 30% glycerol and snap frozen in liquid nitrogen. Diffraction data were collected at the Advanced Photon Source (APS, Argonne, Illinois, USA) and the NSLS2 synchrotron at Brookhaven, New York, USA (Table 1).

### *In vitro* pulldowns and isothermal titration calorimetry

*In vitro* pulldowns were conducted using hexahistidine tagged peptides of interacting proteins as bait, as described in dx.doi.org/10.17504/protocols.io.dm6gpqk91lzp/v2. If not stated otherwise 10μM of bait peptide or the hexahistidine tagged PH domain of OCRL was mixed with 10μM of prey TMEM55B_80-166_ or TMEM55B_80-160_ in 1.5 mL centrifuge tubes with 25 μl Ni^2+^-agarose resin in a final volume of 1ml of binding buffer (20 mM Tris pH 8.0, 300 mM NaCl, 20mM Imidazole, 10 mM β-mercapotoethanol). The reaction mixture was subjected to gentle shaking for 15 minutes. The resin was collected by gentle centrifugation (1,000 rpm) and washed 3 times with 1 ml of the binding buffer. Following release of proteins from the resin with 50μl elution buffer (20 mM Tris-Cl pH 8.0, 300 mM NaCl, 200mM imidazole), samples were subjected to SDS-PAGE and stained with Coomassie Brilliant Blue^26^.

Calorimetry experiments were performed using an ITC200 (Particular Sciences) as described in dx.doi.org/10.17504/protocols.io.j8nlkr7x6v5r/v1. Experiments were performed 3 or more times for each interacting pair. The TBM peptides were dialyzed together with TMEM55B_80-160_ (20 mM) Tris-HCl, 150mM NaCl, 5mM and 1mM DTT, pH 7.5). Protein concentrations were calculated based on their absorbance at 280nm using a ND-1000 NanoDrop spectrophotometer (Thermo Scientific), peptide concentration was determined by adding buffer to a known amount of lyophilized peptide factoring in their purity. The proteins were centrifuged at 13,100 x g for 10 minutes prior to the determination of protein concentration and ITC analyses. 20-45μM of peptide was injected into 200-450μM of TMEM55B. We used Origin 7.0 with the ITC plugin for data processing, and assumed a single binding mode for data fitting.

### Size-exclusion chromatography coupled to multi-angle light scattering

Multi-angle light scattering was performed using a miniDAWN linked to an Optilab rEX refractometer (Wyatt Corp) as described in <u>dx.doi.org/10.17504/protocols.io.3byl41mxolo5/v1</u>. Purified proteins and complexes were adjusted to a concentration of 1-1.5mg/mL, and between 300-400µL was injected into an AKTAbasic chromatography system (Cytiva Life Science). The flow rate was adjusted to 0.6-0.7 mL/min through a Superdex75 10/300 GL column (Amersham, Uppsala, Sweden), which was coupled to the light scattering and refractometer upon elution from the column. Data were recorded and processed with the ASTRA 4.9 software which was also used to calculate the molecular weight of the peaks corresponding to TMEM55B_80-166_ or the complex TMEM55B/RILPL1 peptide. Three independent experiments were conducted for both TMEM55B alone and the complex.

### Generation of HEK293 TMEM55B/A Double Knock-Out Cells by CRISPR-Cas9

A full transcript map of the TMEM55B locus was constructed by combining data from both NCBI (NC_000014.9) and Ensembl (ENSG00000165782). Knockout (KO) guide RNAs were selected to target the exon 2 to ensure complete disruption of all possible transcripts. Three sets of CRISPR-Cas9 guide RNAs were designed to target exon 2 of TMEM55B: a pair targeting exon 2 (Sense A and Anti-sense A); G1, a single guide RNA (3’-GCCCTTAACTAG CCCGGACAG-5’)’ and G2, a single guide RNA (3’-GACTCGGCAGGTGATCATAG-5’). HEK293 cells were co-transfected with 1 μg of each plasmid and 2 μg of PEI mixture supplemented with Opti-MEM. After 24 hr of the transfection, cells were kept in DMEM containing 2 µg/ml of Puromycin for 48 hr. After the recovery, cell pools were analysed for the depletion of TMEM55B expression by immunoblotting and afterwards, single cells were sorted using FACS. Following two to three weeks of recovery, promising clones were verified by PCR, shotgun cloning, and sequencing. In similar way, three sets of CRISPR-Cas9 guide RNAs were designed to target exon 2 of TMEM55A: a pair targeting exon 2 (Sense A and Anti-sense A); G1, a single guide RNA (3’-GCACACACGGCAGTTTATTAC-5’)’ and G2, a single guide RNA (3’-GTGATTAGTGATTGGCACACA-5’). Single sorted HEK293 TMEM55B KO cells were further knocked out and verified for TMEM55A following the same protocol. The above method is also described in <u>dx.doi.org/10.17504/protocols.io.14egnr7q6l5d/v1</u> (RRID: CVCL_F0B6)

### Cell culture, transfection, and lysis

HEK293 cells were purchased from ATCC and maintained in DMEM containing 10% (v/v) FBS, 2 mM L-glutamine, 100 U/ml penicillin, and 100 µg/ml streptomycin. Cells were grown at 37°C temperature with 5% CO_2_ in a humidified atmosphere and regularly tested for mycoplasma contamination. Transient transfections were performed in HEK293 cells 24hr prior to cell lysis using PEI at around 60-70% confluency. Transfections for co-immunoprecipitation assays were done in 10 cm cell culture dishes using 3 µg of Flag-LRRK2 Y1699C or Flag-LRRK2 Y1699C D2017A as indicated, 1 µg of HA-Rab8A Q67L and 2 µg of RILPL1-GFP or Halo-TMEM55B or JIP3-GFP or JIP4-GFP or GFP-OCRL or WDR81-GFP cDNA construct per dish diluted in 1 ml of Opti-MEM and 20 µg of PEI mixture and incubated for 30 min before being added to the media. Cells were lysed in an ice-cold lysis buffer containing 50 mM Tris-HCl, pH 7.5, 1% (v/v) NP-40 alternative or 1% (v/v) Triton-X100, 10% (v/v) glycerol, 150 mM NaCl, 1 mM sodium orthovanadate, 50 mM sodium fluoride, 10 mM sodium β-glycerophosphate, 5 mM sodium pyrophosphate, 0.1 µg/ml microcystin-LR, and 1 tablet of cOmplete Mini (EDTA-free) protease inhibitor (Merck, 11836170001). Protein lysates were clarified by centrifugation at 17,000 x *g* for 10 min and were quantified by Bradford assay. HEK293 cells were sourced ethically, and their research use was in accord with the terms of the informed consents under an IRB approved protocol.

### Co-immunoprecipitation assays

GFP or Halo immunoprecipitation were performed according to the manufacturer’s protocol and as described in dx.doi.org/10.17504/protocols.io.eq2ly7kxqlx9/v1. Briefly, lysates were incubated with either GFP-Trap agarose beads (Chromotek) or HaloLink Resin (Promega) for 1-2 h (20 µl of packed resin/ 1 mg of lysate). Immunoprecipitates were washed three times with wash-buffer (50 mM Tris-HCl pH 7.5, 150 mM NaCl) and then either processed for mass spectrometry experiments or eluted by adding 2x NuPAGE LDS sample buffer for immunoblot analysis. The mixture was then incubated at 95°C for 10 min and the eluent was collected by centrifugation through a 0.22 µm Spin-X column (CLS8161, Sigma). Eluted samples were supplemented with 1% (by volume) β-mercaptoethanol and denatured at 70°C for 10 min before being subjected to immunoblot analysis.

### Quantitative Immunoblotting Analysis

Quantitative immunoblotting analysis was performed according to the protocol described in dx.doi.org/10.17504/protocols.io.bsgrnbv6. Briefly, 10-20 µg of lysate or 25% of the immunoprecipitated samples were loaded onto NuPAGE 4–12% Bis–Tris Midi Gels (Thermo Fisher Scientific, Cat no. WG1402BOX or Cat no. WG1403BOX)) and electrophoresed at constant 120V for 2hr with NuPAGE MOPS SDS running buffer (Thermo Fisher Scientific, Cat no. NP0001-02). At the end of electrophoresis, proteins were electrophoretically transferred onto a nitrocellulose membrane (GE Healthcare, Amersham Protran Supported 0.45µm NC) at constant 90V for 90 min on ice in transfer buffer (48mM Tris and39 mM glycine supplemented with 20% (v/v) methanol). The membranes were blocked with 5% (w/v) skim milk powder dissolved in TBS-T (50mM Tris base, 150mM sodium chloride (NaCl), 0.1% (v/v) Tween 20) at room temperature for 1h. Membranes were washed three times with TBS-T and were incubated in primary antibody overnight at 4°C. Prior to secondary antibody incubation, membranes were washed three times for 10 min each with TBS-T. The membranes were incubated with secondary antibody for 1h at room temperature. Thereafter, membranes were washed with TBS-T five times with a 10min incubation for each wash, and protein bands were acquired via near-infrared fluorescent detection using the Odyssey CLx imaging system and intensities of bands quantified using Image Studio Lite (Version 5.2.5, RRID:SCR_013715).

### Sample preparation, LC-MS/MS, and data analysis for Data-Independent Acquisition (DIA) experiments

The washed Halo immunoprecipitation beads were dissolved in a 100 µl of 50mM TEABC buffer in 6M Urea buffer, mixture was then reduced by adding 5mM DTT and incubated on a Thermomixer at room temperature for 30 min. Followed by the mixture was supplemented with fresh 20 mM (final concentration) Iodoacetamide (stock concentration - 100 mM) was added to the solution and kept in the Thermomixer for 30 min at room temperature with 1250 RPM in dark. 6M urea was then diluted to 1.5M with 50mM TEABC buffer along with 200 ng Trypsin/LysC and incubated for 2h at 30 °C with 1250 RPM. After a brief centrifugation, the supernatant was transferred to a fresh tube and supplemented with another 300 ng Trypsin/LysC and kept in the thermomixer for overnight at 30°C at 1250 RPM. The reaction was quenched by adding final 1% (v/v) Trifluoroacetic acid and peptides were purified using in-house prepared SDB-RP stage-tips. Eluted peptides were vacuum dried and stored at -80°C until LC-MS/MS analysis (Protocol details: https://dx.doi.org/10.17504/protocols.io.kxygxzrokv8j/v1).

#### LC-MS/MS analysis

Lyophilised peptides were dissolved in LC buffer (3% ACN in 0.1% Formic acid) and transferred into LC-vails. The data was acquired on Orbitrap Exploris 480 mass spectrometer in line with Ultimate 3000 nano-RSLC system. Peptides were loaded onto trap column (C18, 5 µm, 100 A°, 100 µ, 2 cm Nano-viper column # 164564, Thermo Scientific) and resolved on 50 cm analytical column (C18, 5 µm, 50 cm, 100 A° Easy nano spray column # ES903, Thermo Scientific) that is directly connected to easy nano-source. LC-MS/MS analysis was performed using variable data independent acquisition (DIA) mode. Complete LC and DIA-MS parameters have been deposited in Zenodo server (https://doi.org/10.5281/zenodo.16881244), and includes the raw files that were processed using DIA-NN 1.8 search algorithm^27^. The database search parameters are also provided in this same Zenodo deposition. The output files from DIA-NN search protein group files were processed using Perseus (version 1.6.15.0) environment ^28^ for data filtering, quantile normalisation, imputation using normal distribution (width = 0.3 and down-shift =1.8). Differential analysis was done between groups using two-sided t-test and multiple testing correction performed with Benjamini Hochberg or Permutation-based FDR correction (5%FDR). The resulting differential analysis files and raw input data further imported into CURTAIN 2.0 for data visualisation and exploration^29^.

**Supplementary Table S1:** Primary Antibodies

| **Antibody Target** | **Source** | **Catalogue Number (RRID)** | **Dilution** |
| --- | --- | --- | --- |
| TMEM55B | Proteintech | 23992-1-AP<br>(AB_2879391) | 1:1000 |
| JIP4/ SPAG9 | Cell Signaling Technology | 5519<br>(AB_10828724) | 1:1000 |
| α-Tubulin | Cell Signaling Technology | 3873<br>(AB_1904178) | 1:10000 |
| Halo | Promega | G9211<br>(AB_2688011) | 1:1000 |
| LRRK2 Total C-terminal | Neuromab | N241A/34<br>(AB_2877351) | 1 μg/ml |
| Phospho-Rab8A (Pan-Thr-specific) | Abcam | ab230260<br>(AB_2814988) | 1:1000 |
| HA | Roche | 3F10<br>(AB_2314622) | 1:1000 |
| GFP | ChromoTek | 3H9<br>(AB_10773374) | 1:1000 |
| RILPL1 | Abcam | Ab302492<br>(AB_2936945) | 1:1000 |
| TMEM55A | MRC PPU Reagents and Services, University of Dundee | DA241<br>(AB_2936451) | 1 μg/ml |

**Supplementary Table S2:** Secondary Antibodies

| **Antibody Target** | **Company** | **Catalogue number (RRID)** | **Dilution** |
| --- | --- | --- | --- |
| IRDye 800CW Goat anti-Rabbit IgG | LI-COR | 926-32211<br>(AB_621843) | 1:25,000 |
| IRDye 680CW Goat anti-Mouse IgG | LI-COR | 926-68070<br>(AB_10956588) | 1:25,000 |
| IRDye 680CW Goat anti-Rat IgG | LI-COR | 926-68074<br>(AB_10956736) | 1:25,000 |
| IRDye 800CW Donkey anti-Rabbit IgG | LI-COR | 926-32213<br>(AB_621848) | 1:25,000 |
| IRDye 680LT Donkey anti-Mouse IgG | LI-COR | 926-68 022<br>(AB_10715072) | 1:25,000 |
| IRDye 680LT Donkey anti-Goat IgG | LI-COR | 926-68 024<br>(AB_10706168) | 1:25,000 |

---

**Figure S1.**
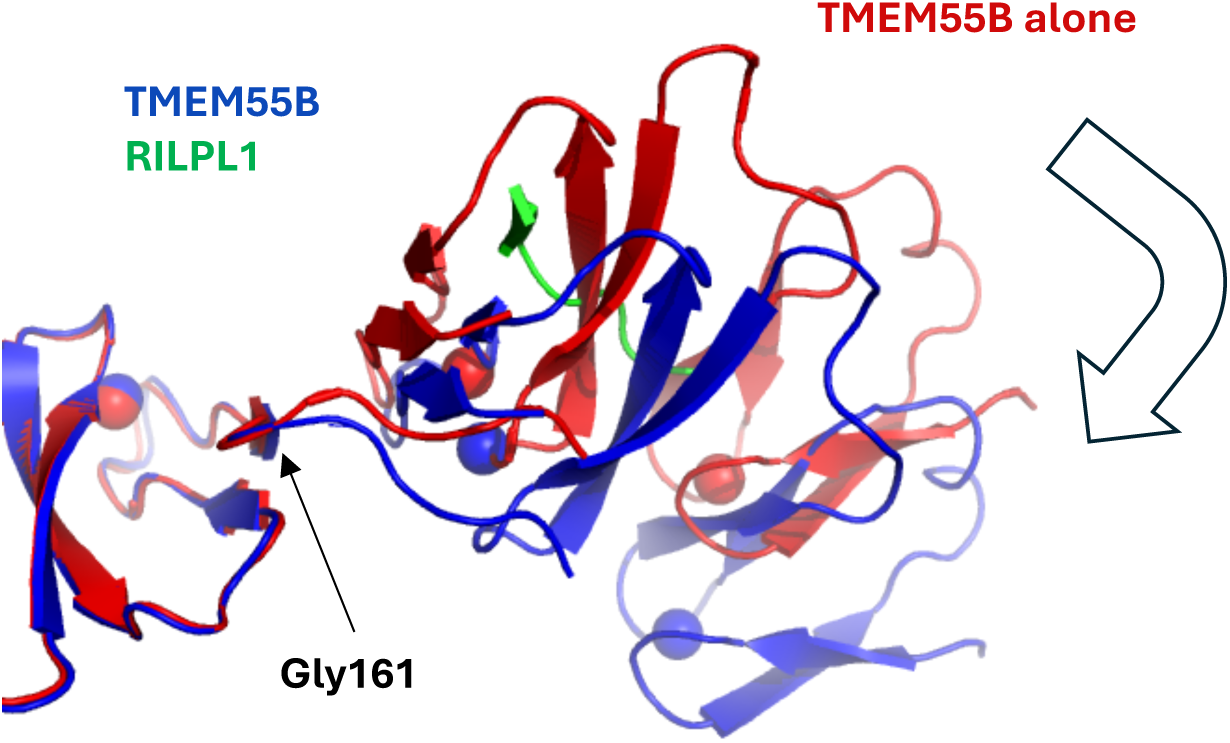
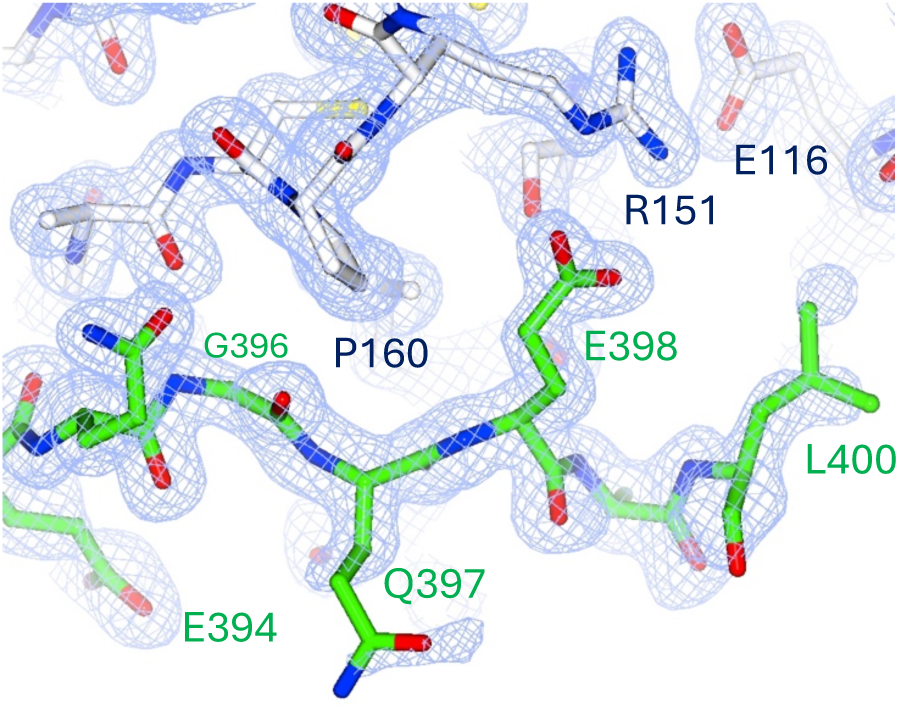
**A.** Ribbon models of TMEM 55B alone and in complex with RILPL1. The asymmetric unit is a dimer in both crystals. Molecule B has been superposed (left), and molecule A shows a relative rotation of approximately 30^°^. The hinge for this difference is Gly161, whose (angle rotates by 37^°^. The list of (((((are below: TMEM55B alone = L160 (−87.4^°^, 9.8^°^), G161 (−176.1^°^, 158.9^°^), P162 (−67.5^°^, 157.5^°^); TMEM55B complex = L160 (−106.5 ^°^, 26.3^°^), G161 (−139^°^, 164.8^°^), P162 (−80.7 ^°^, 164.7 **B.** A section of the electron density around the TBM motif from the crystal structure of TMEM55B/RILPL1 complex. The RILPL1 peptide is green, and the TMEM55B is grey. The 2Fo-Fc map is contoured at 1α density.

**Figure S2:**
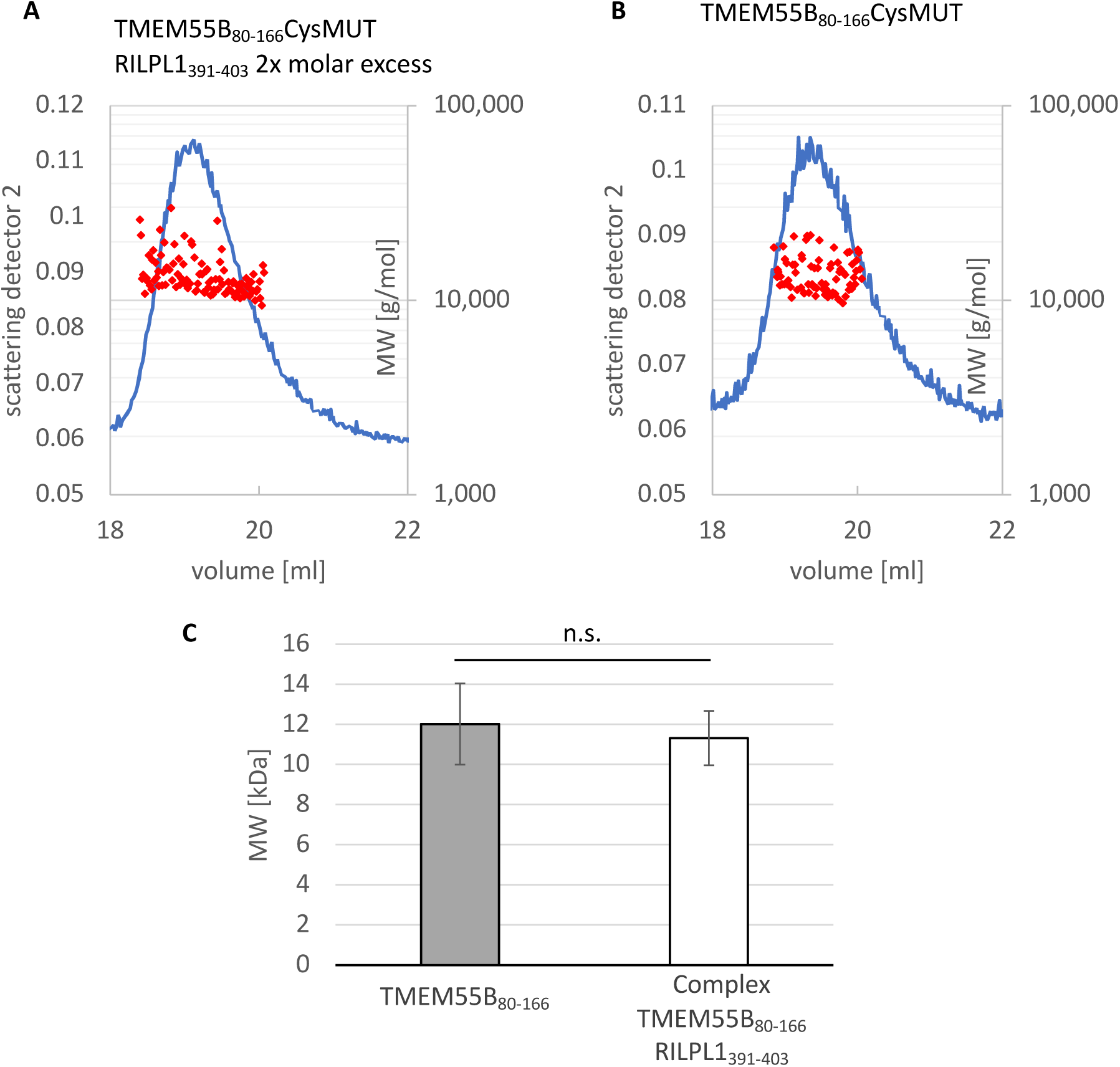
Size exclusion chromatography coupled to light scattering was performed to compare TMEM55B_80-166_2CysMUT alone with a sample pre-incubated with peptide RILPL1_391-403_. Samples were injected onto a Superdex75 10/300 column coupled to the Wyatt systems described in Methods. Absolute mass measurements were calculated at 1 second intervals. **(A)** and **(B)** show representative runs of TMEM55B/RILPL1 and TMEM55B alone, respectively. **(C)** Bar plot of the calculated molecular weights from 3 independent experiments. For each independent experiment, the value of the molecular weight was calculated as the average over the peaks (red dots in **A** and **B**).

**Figure S3:**
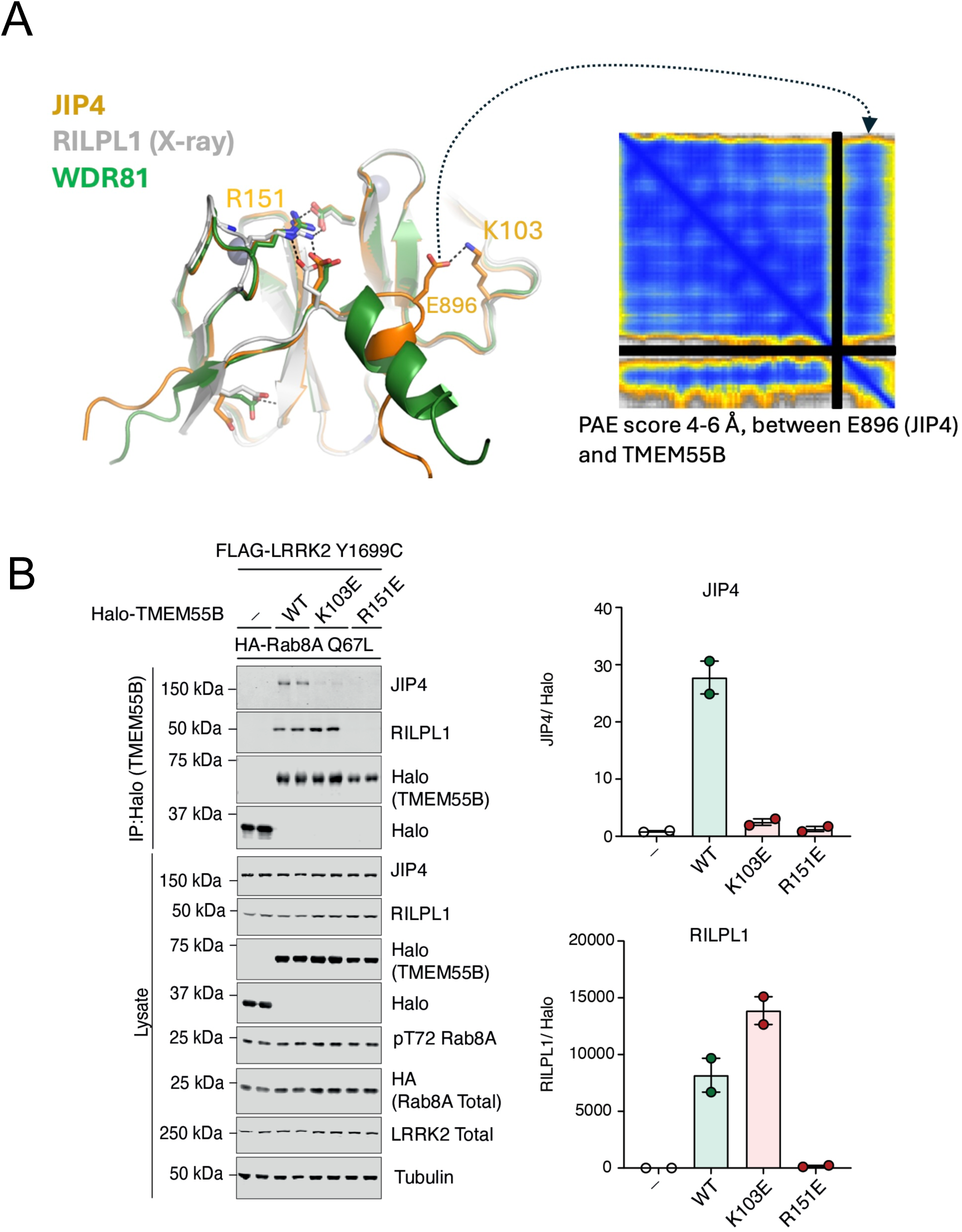
(A) AlphaFold3 predictions of TMEM55B in complex with JIP4 and WDR81. Peptide predictions are aligned onto the X-ray structure of RILPL1 peptide. The panel on the right shows a graphic of PAE scores from the JIP4/TMEM55B model, visualized using ChimeraX (ref). JIP3 was not included in the alignment because AlphaFold3 failed to yield a reasonable model. (B) HEK293 TMEM55B/A double KO cells were transiently transfected with HA-Rab8(Q67L) and FLAG-LRRK2(Y1699C, kinase-active mutant), in the presence of WT, K130E and R151E variants of Halo-TMEM55B. Halo-IP was performed and analyzed by quantitative immunoblot analysis using the LiCOR Odyssey CLx Western blot imaging system and indicated antibodies. Quantitation of immunoblotting data are dhown as mean +/-SEM (ImageStudio Lite version 5.2.5).

**Figure S4:**
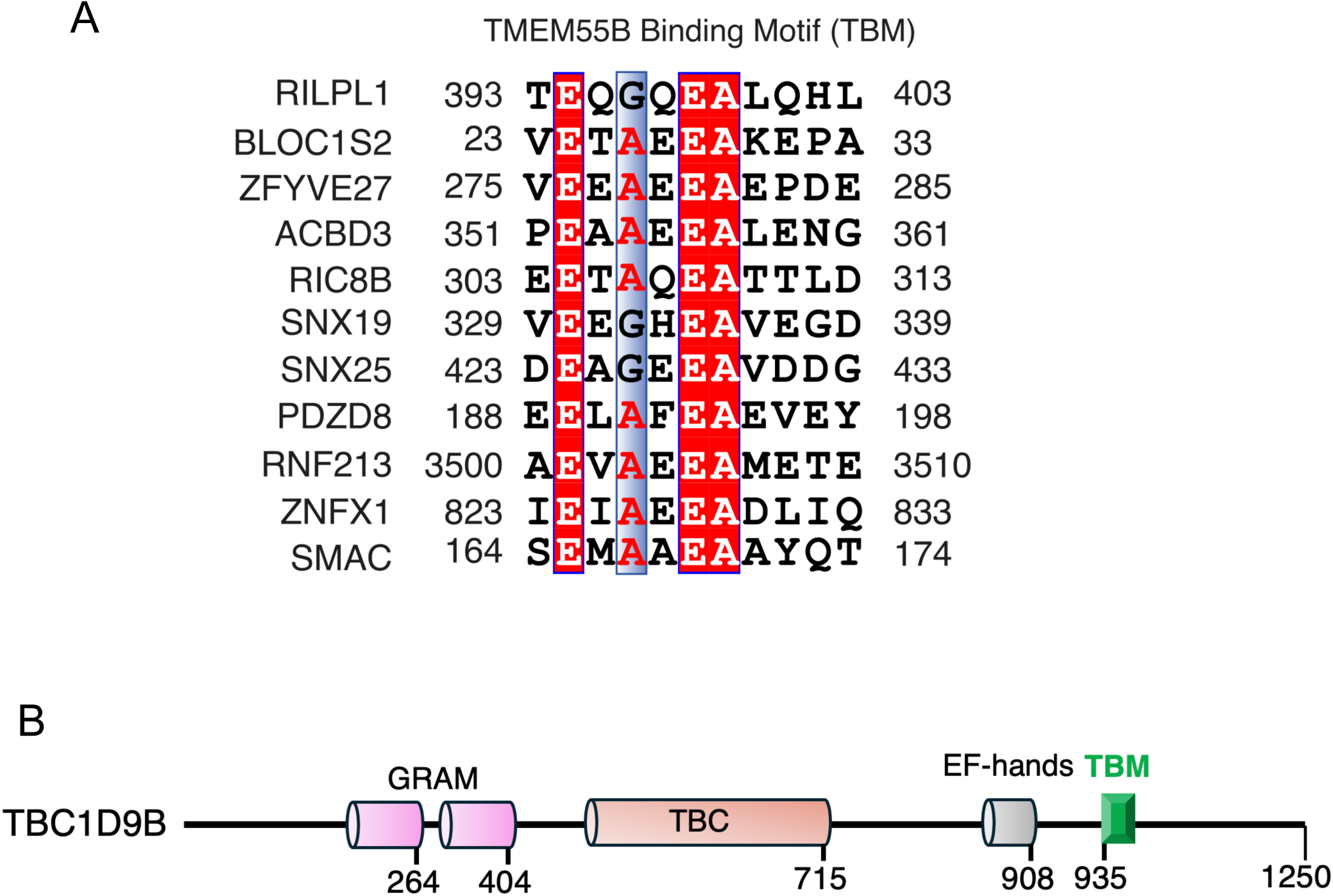
(A) Potential TBMs from interactors identified by mass spectrometry. These proteins have not been verified by independent experiments. (B) Domain organization of TBC1D9B with location of the TBM indicated in green.

**Figure S5:**
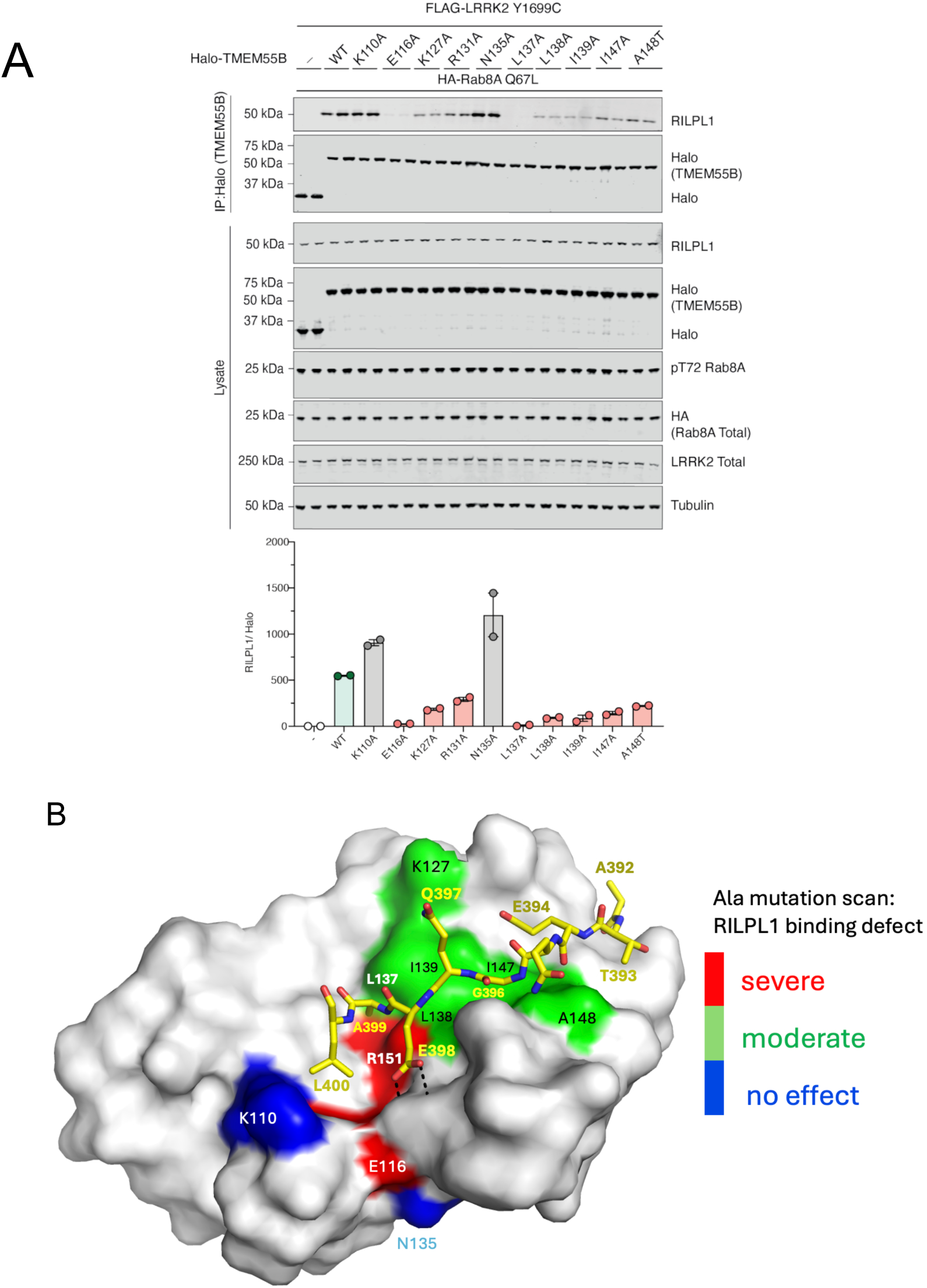
**(A)** HEK293 TMEM55B/A double knockout cells were transiently transfected with HA-Rab8a(Q67L GTP-bound mutant) and FLAG-LRRK2(Y1699C kinase-active mutant) either in the presence of Halo-TMEM55B WT or ten different mutants (K110A, E116A, K127A, R131A, N135A, L137A, L138A, I139A, I147A, A148T) at the TMEM55B interface. Halo immunoprecipitation was performed and analyzed by quantitative immunoblot analysis using the LI-COR Odyssey CLx Western blot imaging system and indicated antibodies. Quantitation of immunoblotting data (performed using ImageStudioLite software version 5.2.5, RRID:SCR_013715) is shown as mean ±SEM. **(B)** Surface map of TMEM55B in complex with stick model of RILPL1 peptide (yellow). Residues on TMEM55B surface that were mutated to alanine are colored red (negatively charged), blue (positive), cyan (polar) and green for non-polar side chains. The key electrostatic interaction interaction between E398^RL^^1^ and R151™ is emphasized with dashed black line reaching out to the grey surface contours of arginine.

## Notes

https://doi.org/10.5281/zenodo.16883734

https://doi.org/10.5281/zenodo.16883557).

https://doi.org/10.5281/zenodo.16814518

https://doi.org/10.5281/zenodo.16814483

https://doi.org/10.5281/zenodo.16814276

https://doi.org/10.5281/zenodo.16881244

### Competing Interest Statement

The authors have declared no competing interest.

